# A Novel, Uniquely Efficacious Type of CFTR Corrector with Complementary Mode of Action

**DOI:** 10.1101/2023.08.15.552928

**Authors:** Valentina Marchesin, Lucile Monnier, Peter Blattmann, Florent Chevillard, Christine Kuntz, Camille Forny, Judith Kamper, Rolf Studer, Alexandre Bossu, Eric A. Ertel, Oliver Nayler, Christine Brotschi, Jodi T. Williams, John Gatfield

## Abstract

Three distinct pharmacological corrector types (I, II, III) with different binding sites and additive behaviour only partially rescue the F508del-CFTR folding and trafficking defect observed in cystic fibrosis. Here, we describe novel, uniquely effective, macrocyclic CFTR correctors that were additive to the known corrector types, thus exerting a new, complementary “type-IV” corrector mechanism. Macrocycles achieved wildtype-like folding efficiency of F508del-CFTR at the endoplasmic reticulum and normalized CFTR currents in reconstituted patient-derived bronchial epithelium. Using photo-activatable macrocycles, docking studies and site-directed mutagenesis a highly probable binding site and pose for type-IV correctors was identified in a cavity between lasso helix-1 (Lh1) and transmembrane helix-1 of membrane spanning domain-1 (MSD1), distinct from the known corrector binding sites. Since only F508del-CFTR fragments spanning from Lh1 until MSD2 responded to type-IV correctors, these likely promote co-translational assembly of Lh1, MSD1, and MSD2. Remarkably, previously corrector-resistant CFTR folding mutations were also robustly rescued, suggesting substantial therapeutic potential for this novel type-IV corrector mechanism.

**Teaser:** A novel type of macrocyclic CFTR corrector with new binding site, complementary mode of action and unique folding / trafficking efficacy is described.

## INTRODUCTION

Cystic fibrosis is a common life-threatening genetic disorder caused by mutations in the Cystic Fibrosis Transmembrane Conductance Regulator (CFTR) gene [1]. CFTR is a chloride and bicarbonate channel expressed on the surface of secretory epithelia. Its absence or dysfunction compromises mucus hydration and fluidity leading to mucus accumulation and chronic inflammation in multiple organs and their progressive functional impairment, with the most severe, ultimately lethal effects in the lung [2, 3]. In recent decades, early diagnosis and symptomatic treatments have improved CF patientś quality of life and extended their life expectancy [4]. In the last 10 years, CFTR-targeted therapies (CFTR modulators) have further improved symptoms and quality of life, demonstrating that addressing the molecular defect in CF is an effective strategy and warrants further research [3].

CFTR is a 1480 amino acid transmembrane protein of the ABC transporter family. It consists of two membrane spanning domains (MSD1, MSD2), two nucleotide binding domains (NBD1, NBD2) and a regulatory domain whose phosphorylation initiates channel opening [5, 6]. More than 700 disease-causing mutations in the CFTR gene are known to date (http://cftr2.org). The most prevalent mutation (∼80% of patients) is a F508 deletion in NBD1 [3]11 which interferes with CFTR folding at the ER by reducing NBD1 thermal stability and damaging the NBD1-MSD2 interface [7, 8]. Misfolded CFTR is prematurely degraded by the ER-associated degradation machinery, thus strongly reducing CFTR trafficking to the plasma membrane (PM). The discovery of pharmaco-chaperones (“CFTR correctors”), small molecules that bind to distinct sites on F508del-CFTR and promote its correct folding and trafficking, has revolutionized CF treatment [3, 9]. Based on their additive behavior in rescuing folding-deficient CFTR and based on their different binding sites, CFTR correctors can be grouped into three types [7, 10, 11] with a developing nomenclature. Type-I correctors (lumacaftor, LUM; tezacaftor, TEZ; galicaftor, GAL; all with highly similar structures) bind to a recently defined binding site within MSD1 and stabilize this domain [12–15] with allosteric effects on overall F508del-CFTR folding. Type-I correctors show low efficacy in vitro and in humans as monotherapy [16–19]. Non-clinical type-II correctors (Corrector4a; C4a) are thought to bind to the NBD2 domain to promote F508del-CFTR folding and trafficking [20], although their binding site has not been identified yet. Finally, the most recently described corrector type, represented by elexacaftor (ELX) and bamocaftor (BAM) that have very similar chemical structures [16, 17], has been categorized as “type-III” and was initially thought to bind to and stabilize NBD1 [7, 10, 11]. However, the recently discovered binding site at an interface between the N-terminal lasso domain and MSD2-transmembrane helices (TM)-10/11, rather suggests that these correctors assist in the assembly of MSD1 and MSD2 [21]. The classification of corrector types (type-I, -II, -III) that is based on their chronological appearance, is followed throughout this manuscript. Type-III correctors can have higher efficacies than type-I and type-II correctors and, when combined with type-I correctors, F508del-CFTR folding and trafficking is rescued to ∼50 % of normal (TEZ+ELX) [10, 11]. The most recently approved therapy, Trikafta®, combines type-III corrector ELX and type-I corrector TEZ together with a channel opener ivacaftor (IVA), and causes a substantial improvement in lung function in F508del patients [16, 17, 22, 23]. The clinical efficacy of Trikafta is superior to the previous combinations of LUM and IVA (Orkambi®) or TEZ and IVA (Symdeko®), suggesting that co-treatment with multiple correctors of different types improves correct trafficking of folding-deficient CFTR [22, 23], through co-operative pharmaco-chaperoning via the different CFTR binding sites [11]. Considering the less than full rescue even after combining the known corrector types, the identification of highly effective corrector types, with new binding sites and complementary modes of action, is warranted.

Here, we describe macrocyclic CFTR correctors from our drug discovery program, which achieved wildtype-like folding efficiency of F508del-CFTR through a novel type-IV correction mechanism. The corrected F508del-CFTR chloride current in reconstituted CF patient-derived bronchial epithelium matched currents of non-CF controls. Using photo-cross-linkable macrocyclic probes, CFTR peptide mapping, site-directed mutagenesis and molecular modeling, a cavity between CFTR lasso helix-1 (Lh1) and MSD1-TM1/2 was identified as the most likely macrocycle binding site. This site is distinct from reported binding sites of type-I, -II and -III correctors, which is consistent with the additive behaviour with known correctors. We propose that type-IV correctors co-translationally stabilize Lh1-MSD1-MSD2 interactions, thus drastically increasing the folding efficiency of mutated CFTR. Accordingly, type-IV correction also rescued many other CFTR folding mutations including ones resistant to TEZ+ELX treatment, demonstrating promising therapeutic potential for this novel corrector type.

## RESULTS

### Type-IV correctors robustly restore F508del-CFTR trafficking and behave additively with type-I, type-II, and type-III correctors

To identify CFTR correctors with novel chemotypes and a complementary mode of action, we performed a high-throughput screening (HTS) campaign on the Idorsia compound library to discover compounds that increased F508del-CFTR surface expression after overnight treatment and behaved additively with the known corrector types. We used U2OS cells in which cell surface-residing F508del-CFTR was quantified via enzyme fragment complementation (pharmacotrafficking assay). The 17-membered macrocycles depicted in figure 1A represent a selection of mechanistically novel CFTR correctors derived from our drug discovery program which started with one of the HTS hits. These macrocycles were compared with type-I, -II and -III correctors in the pharmacotrafficking assay. All correctors increased F508del-CFTR surface expression with different potencies and maximal intrinsic efficacies (Figure 1B, Figure S1A). The type-I correctors (TEZ, LUM, GAL) and the type-II corrector C4a increased F508del-CFTR surface expression maximally 4- to 5-fold over baseline whereas the type-III correctors (ELX, BAM) increased surface expression maximally 7- to 9-fold. In contrast, the selected macrocycles achieved maximal surface expression of 7-fold (IDOR-1) to 14-, 15- and 18-fold (IDOR-2, -3, -4), respectively. Macrocycle potency ranged between EC_50_=560 nM (IDOR-1) and EC_50_=18 nM (IDOR-4), which was up to tenfold more potent than either TEZ, LUM, BAM or ELX (Figure 1B, Figure S1A). ELX, titrated onto a maximally effective concentration of TEZ, reached in combination an E_max_ of 10-fold over baseline demonstrating the expected additivity for type-I and type-III correctors. Corrector activities were fully confirmed by classical anti-CFTR immunoblotting (C-band) in the same samples (Figure S1B-C). Furthermore, none of the CFTR correctors promoted trafficking of misfolded adrenergic-β2 receptor ADBR2^W158A^ (Figure S1D), which excludes unspecific effects of correctors on ER quality control.

**Fig. 1.**
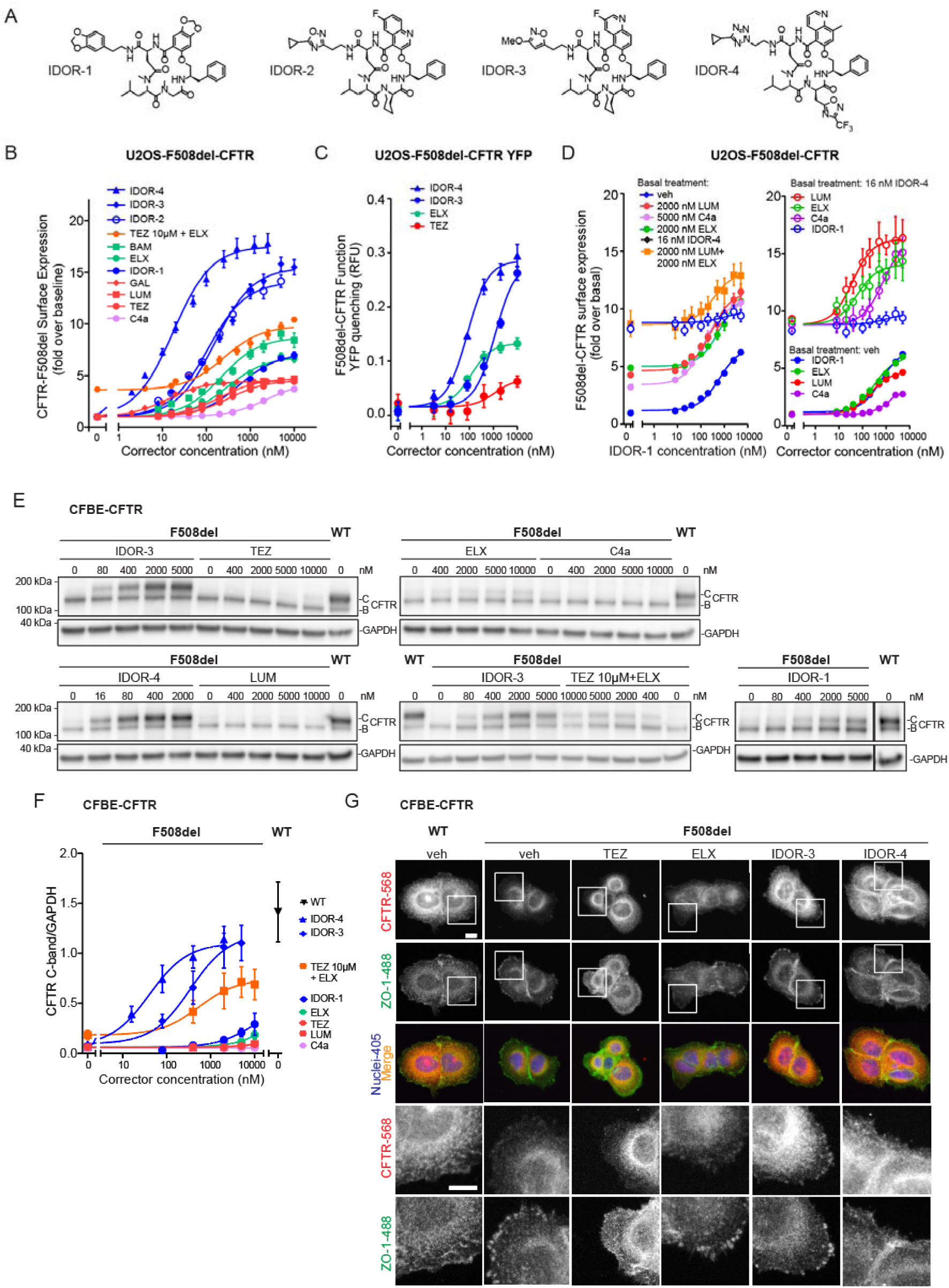
Type-IV correctors restore F508del-CFTR trafficking to the cell surface and behave additively with type-I, type-II, and type-III correctors. (A) Structures of the macrocyclic type-IV correctors used in this study. (B) F508del-CFTR cell surface expression in U2OS cells after overnight treatment with different concentrations of the indicated correctors (n≥4). (C) F508del-CFTR function (YFP quenching assay) after 24 h treatment of U2OS-F508del-CFTR-YFP cells with different concentrations of the indicated correctors in the acute presence of 0.1 µM forskolin and 2 nM IVA (n=2). (D) Left panel: F508del-CFTR cell surface expression in U2OS cells after overnight treatment with different concentrations of IDOR-1 (n=3) in presence of vehicle or a maximally effective concentration of type-I corrector (LUM), type-II corrector (C4a), type-III corrector (ELX), type-I + type-III correctors (LUM+ELX) or a sub-maximally effective concentration of IDOR-4. Right panel: F508del-CFTR cell surface expression in U2OS cells after overnight treatment with different concentrations of type-I corrector (LUM), type-II corrector (C4a), type-III corrector (ELX) or type-IV corrector (IDOR-1) (n=3) in presence of vehicle or a sub-maximally effective concentration of type-IV corrector (IDOR-4). (E) Immunoblot analysis of CFBE41o-cells expressing F508del-CFTR or WT-CFTR treated for 24 h with different concentrations of the indicated correctors and probed for CFTR and GAPDH. Representative images of 3 experiments. (F) Quantification of the CFTR C-band intensities in (E) normalized for GAPDH (n=3). (G) Localization of CFTR (red) and ZO-1 (green) by immunofluorescence confocal microscopy in CFBE41o-cells treated with either DMSO, 2 µM of IDOR-3, 2 µM of IDOR-4, or 10 µM of the other indicated correctors for 24 h. Nuclei appear blue. Scale bar: 10 µm. Data in (B), (C), (D), (F) are means ± SEM of the indicated number (n) of independent experiments. See also Figure S1 and Figure S2.

F508del-CFTR function was assessed in yellow fluorescent protein (YFP) quenching assays [24], in the presence of forskolin and the CFTR potentiator IVA (2 nM, ∼EC_80_): overnight treatment of U2OS-F508del-CFTR cells expressing a halide-sensitive YFP with IDOR-3 or IDOR-4 strongly increased F508del-CFTR function (Figure 1C). ELX and TEZ were ∼2.5- to 6-fold less efficacious, confirming the efficacy ranking of the pharmacotrafficking assay. Responses were blocked by the CFTR inhibitor I-172 (Figure S1F) and activity was lower in absence of IVA (Figure S1E). Some CFTR correctors possess acute inhibitory (e.g., GLPG-2737 [25]), or co-potentiating effects (ELX, [21, 26, 27]) on CFTR currents. In HEK293 cells expressing the gating mutant G551D-CFTR, incubation (10 min) with IDOR-3 or IDOR-4 caused co-potentiation of IVA activity, similar to the effect of ELX, while a corrector from the company Galapagos (WO2017060874, example#331) displayed an acute inhibitory effect (Figure S1H). These acute effects on a CFTR mutant that does not have a trafficking deficit, suggest a direct binding of macrocycles to CFTR without inhibiting its activity, similar to the co-potentiator effect of ELX [21].

To analyze whether these macrocycles represent a new corrector type with a mechanism complementary to existing correctors, we used moderately active IDOR-1 to be titrated onto maximally effective concentrations of selected reference correctors (Figure 1D left panel). Overnight treatment with IDOR-1 increased F508del-CFTR surface expression when used alone, and IDOR-1 was fully additive over its whole concentration range on top of type-I corrector LUM, type-II corrector C4a, or type-III corrector ELX as demonstrated by the strictly parallel upward shift of the IDOR-1 CRCs in the combinations. Only when applied on top of another macrocycle (IDOR-4), IDOR-1 did not achieve further correction (Figure 1D left panel).

A similar lack of additivity was seen when type-I corrector LUM was applied on top of another type-I corrector TEZ (Figure S2A) or when the type-III corrector ELX was titrated onto the type-III corrector BAM (Figure S2B). Also, IDOR-1 was additive to the combination of type-I corrector LUM and type-III corrector ELX (Figure 1D left panel). The more effective IDOR-3 and IDOR-4 (Figure S2C) showed analogous additive behavior at the lower concentrations before reaching the plateau, which likely represents the maximally possible correction level in this assay. In a complementary approach, reference correctors LUM, C4a and ELX were titrated onto a highly effective concentration of IDOR-4. Also in this format, full additivity or even synergy was observed for the type-I, -II and -III correctors while IDOR-1 displayed no additivity with IDOR-4 (Figure 1D right panel). In conclusion, macrocycles displayed a clearly additive efficacy on top of the known corrector types and should therefore possess a new, complementary mode of action and a distinct CFTR binding site.

Next, we performed anti-CFTR immuno-blotting in cystic fibrosis bronchial epithelial cell lines (CFBE41o-) expressing either F508del-CFTR or WT-CFTR [28]. Both cell lines express similar mRNA levels of F508del-CFTR or WT-CFTR, and neither macrocycles nor ELX/TEZ affected these levels, which excludes effects on transcription (Figure S2D). Also, macrocycles did not block the proteasome as deduced from unchanged ubiquitinated protein levels (Figure S2E-F), excluding this unspecific mechanism. IDOR-3 and IDOR-4 concentration-dependently induced strong F508del-CFTR C band accumulation (Figure 1E) reaching ∼75 % of the levels observed in WT-CFTR-expressing cells, whereas TEZ, LUM and ELX showed lower efficacies (Figure 1F) as already observed in the U2OS system. Again, ELX titrated onto a maximally effective concentration of TEZ behaved additively as expected (Figure 1F). Interestingly, IDOR-3 and IDOR-4 lead to a ∼2-fold increase in WT-CFTR C-band (Figure S2G-H) suggesting that even the complex folding of WT-CFTR can profit from highly effective pharmaco-chaperones.

Increased F508del-CFTR C-band expression in CFBE cells reflected trafficking to the cell surface as shown by immunofluorescence microscopy. The level of F508del-CFTR located in plasma membrane ruffles, leading edges, or cell-cell contacts, and co-localizing with the plasma membrane protein ZO-1 was considerably higher in cells incubated with IDOR-3 or IDOR-4 as compared to cells treated with TEZ, ELX, or vehicle (which displayed mainly intracellular CFTR staining reminiscent of the ER), and closely resembled the localization of WT-CFTR (Figure 1G, Figure S2I).

In summary, we have identified a novel type of CFTR corrector characterized by unprecedented efficacy in promoting folding, trafficking, and function of F508del-CFTR and by additivity with the existing corrector types. We classify these macrocycles with complementary mechanism as type-IV correctors.

### Type-IV correctors rescue F508del-CFTR trafficking and function in reconstituted cystic fibrosis bronchial epithelium

Type-IV correctors were analyzed in reconstituted human bronchial epithelia derived from biopsies of CF and non-CF donors and cultivated at air-liquid-interface. In tissues from three CF patients (all F508del-F508del), IDOR-3 and IDOR-4 concentration-dependently increased CFTR C-band formation (Figure 2A-B, Figure S3A) reaching levels (CF patients #2, #3) matching the four non-CF donor controls (Figure 2C). The effects of TEZ and ELX were inferior and difficult to quantify, as reported previously [17].

**Fig. 2.**
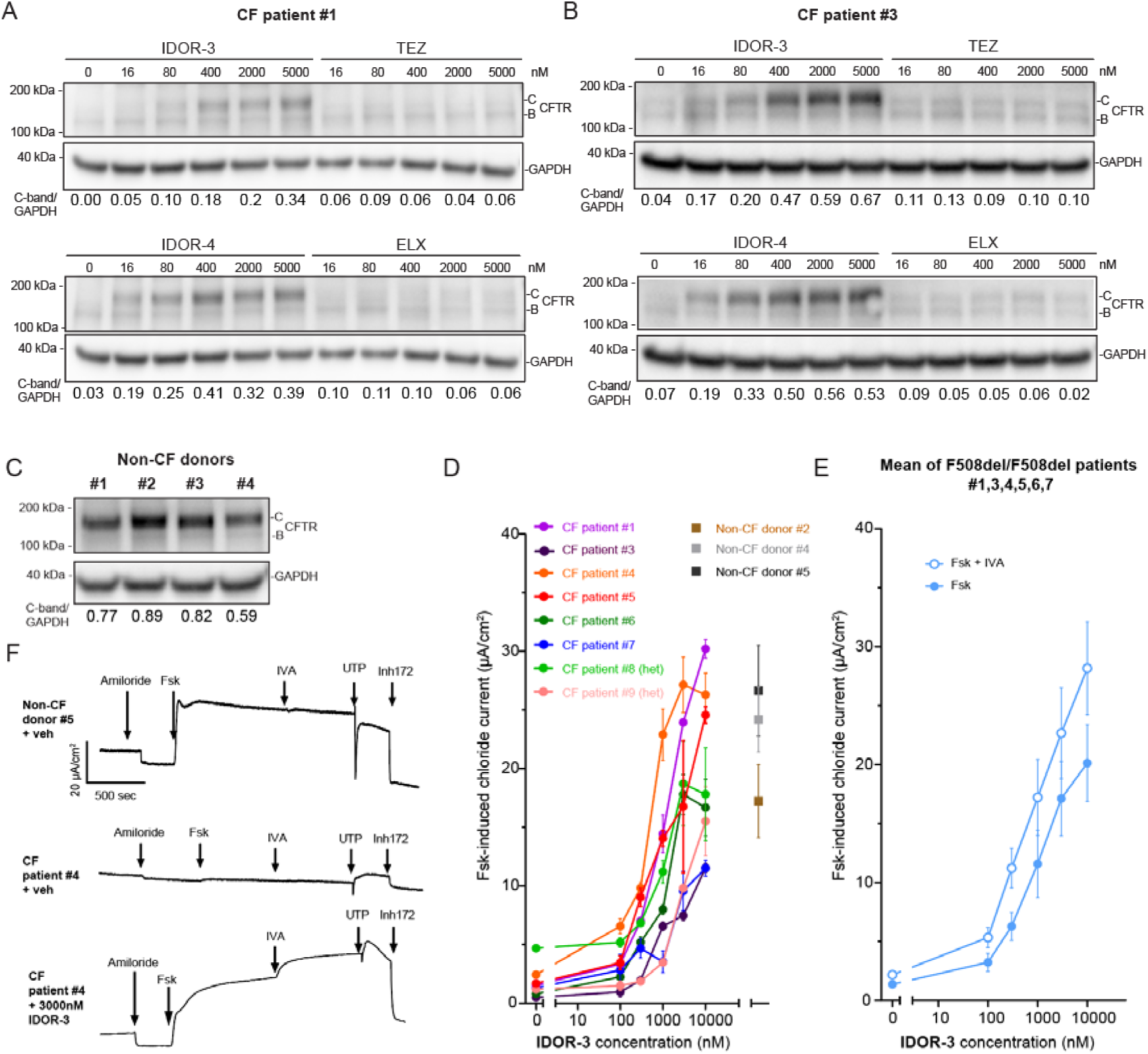
Type-IV correctors rescue F508del-CFTR trafficking and function in reconstituted cystic fibrosis bronchial epithelium. **(A-C)** CFTR expression (immunoblotting) in reconstituted human bronchial epithelium of two F508del-CFTR homozygous CF patients and four non-CF donors after a 24-h treatment with the indicated correctors. Representative images of 2 experiments. The CFTR C-band/GAPDH intensity ratio is represented below every lane. **(D)** Transepithelial forskolin-induced short-circuit currents (Ussing chamber) in reconstituted epithelia of eight CF patients (six F/F, two F/MF) treated overnight with the indicated correctors or vehicle; additionally, tissues from three non-CF donors were treated overnight with vehicle (n≥2 measurement per patient/per concentration). **(E)** Mean transepithelial forskolin-induced short-circuit currents for the six F/F patients in (D) with or without acute IVA addition (1 µM). **(F)** Representative examples of raw current traces recorded during a baseline period followed by sequential addition of amiloride, forskolin, IVA, UTP and Inh172 taken from the experiments described in (D, E). Data in (D), (E) are means ± SEM. See also Figure S3.

To measure recovery of CFTR function, trans-epithelial chloride currents were measured in Ussing chambers using reconstituted tissue from some of the same and some additional donors (6 F508del/F508del, 2 F508del/Minimal Function, 3 non-CF). IDOR-3 concentration-dependently restored forskolin-induced chloride currents to densities observed in the non-CF tissues with some potency and efficacy differences between donors (Figure 2D), even in the absence of a potentiator. Acute IVA addition resulted in a leftward shift of the IDOR-3 concentration-response curve (Figure 2E-F, Figure S3B for individual patients). IDOR-4 was tested in reconstituted epithelia of three CF donors with results similar to IDOR-3, but further improved potency (CF patient #1, #4, #5, Figure S3B). TEZ and ELX, tested on CF patients #3 and #6, showed considerably lower efficacies compared to IDOR-3 (Figure S3B). Altogether these data using primary reconstituted CF bronchial epithelia confirm the high efficacy of type-IV correctors in promoting F508del-CFTR trafficking and function.

### Type-IV correctors restore F508del-CFTR folding efficiency in the endoplasmic reticulum beyond wildtype levels

Next, we analyzed the correction kinetics in CFBE cells expressing F508del-CFTR (Figure 3A-B). IDOR-3, IDOR-4, and ELX induced C-band increases, reaching steady state after ∼8 h. Correction by IDOR-3 and ELX was reversible upon wash-out with a half-life of ∼4-6 h. The effect of IDOR-4 was less reversible possibly due to incomplete wash-out. Thus, the continued presence of correctors is needed to yield stable F508del-CFTR levels.

**Fig. 3.**
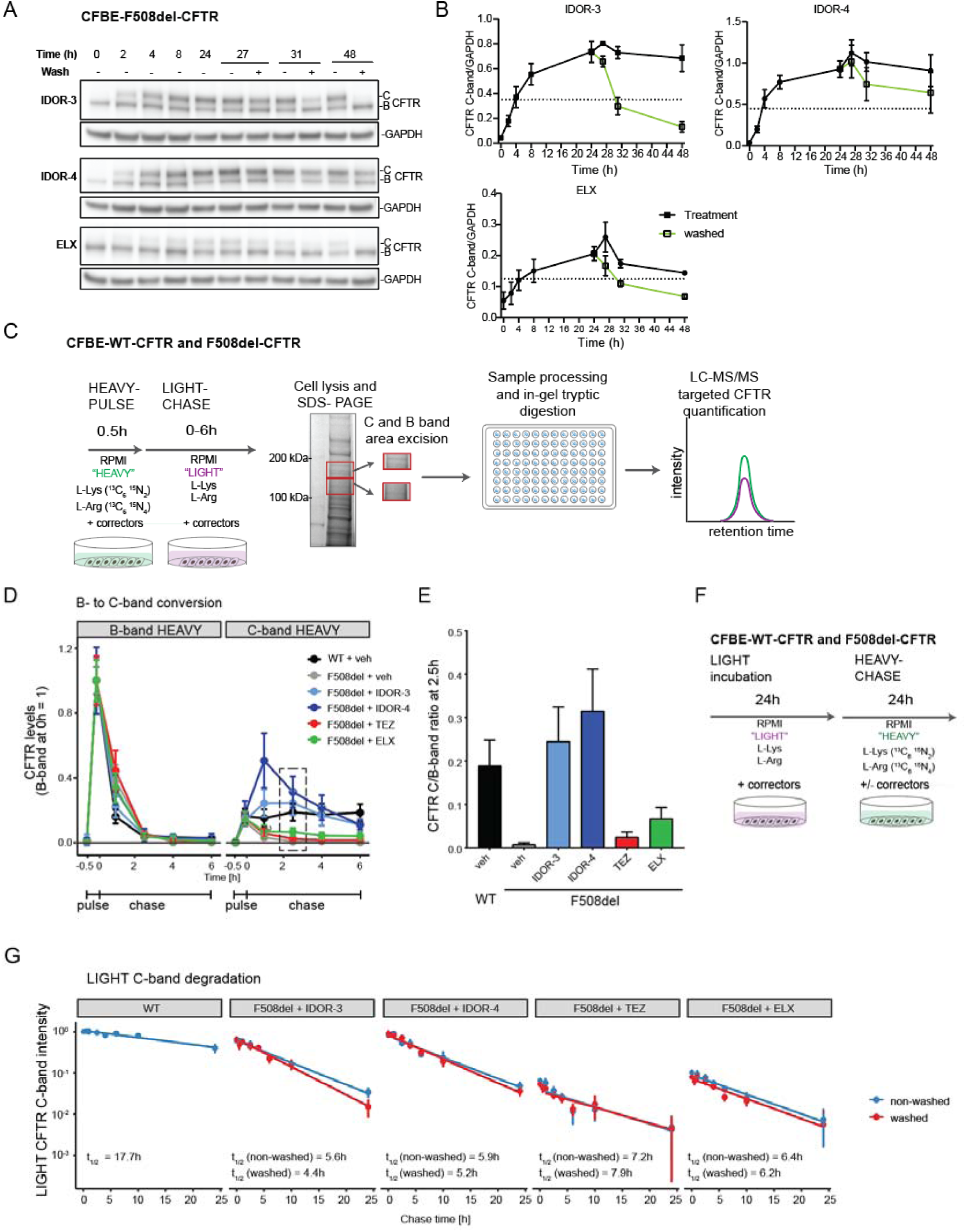
Type-IV correctors restore F508del-CFTR folding efficiency in the endoplasmic reticulum beyond wildtype levels. **(A)** F508del-CFTR correction kinetics (immunoblotting) in CFBE41o-cells expressing F508del-CFTR treated with 2 µM of IDOR-3 or IDOR-4 or 10 µM of ELX for the indicated durations including a corrector wash-out at 24 h. Representative images of 2 experiments. **(B)** Quantification of F508del-CFTR C-band intensities in (A) normalized for GAPDH (n=2). Dashed line represents the midpoint between minimum value at 0 h and maximum value at 24 h. **(C)** Workflow to determine the CFTR folding efficiency at the ER using SILAC-based heavy-pulse-light-chase assays in CFBE41o-cells expressing WT- or F508del-CFTR. **(D)** Average normalized peptide intensities of heavy CFTR from B-bands (left) and C-bands (right) of cells labeled with a 30 min pulse of heavy amino acids and chased with light amino acids. Cells were treated with either DMSO (n=5), 2 µM IDOR-3 (n=4), 2 µM IDOR-4 (n=2), 10 µM TEZ (n=3), or 10 µM ELX (n=3) during both pulse and chase periods. Per treatment, peptide intensities at the different time points were normalized to the corresponding maximal heavy B-band intensity occurring directly after the pulse. **(E)** Folding efficiency determined as average intensity ratio of heavy C-band (2.5 h chase, dashed square in (D) and heavy B-band (directly after pulse) for the different treatments shown in (D). **(F)** Workflow to determine CFTR stability after folding. **(G)** CFTR C-band half-life with and without corrector wash-out: cells were incubated for 24 h with light amino acids in the presence of either DMSO (n=4), 2 µM IDOR-3 (n=2), 2 µM IDOR-4 (n=2), 10 µM TEZ (n=2), or 10 µM ELX (n=2) and then chased with heavy amino acids. Correctors were either kept during the chase period (blue lines) or washed-out before the chase (red lines). Half-life was determined by linear regression of the logarithmic data. Data in (B), (D), (E) and (G) are means ± SEM of the indicated number of independent experiments. See also Figure S4.

To analyse if type-IV correctors work via increasing CFTR folding efficiency in the ER [7, 11, 29] we developed a metabolic pulse-chase assay using stable isotope labeling of amino acids in cell culture (SILAC) [30]. Proteins in CFBE cells were labeled using a 30 min pulse of medium containing heavy isotopologues of arginine and lysine, followed by a chase in medium containing light amino acids (Figure 3C). Correctors at maximally efficacious concentrations were present during pulse and chase phases. Cells were then lysed at various time points and proteins were separated by SDS-PAGE. Gel regions containing CFTR B- and C-bands were excised separately, trypsin digested and analyzed by high-resolution mass spectrometry (Figure 3C, Figure S4A-C). The pulse resulted in heavy-labeling of ∼30% (WT-CFTR) or ∼50% (F508del-CFTR) of the immature CFTR B-pool (Figure S4D). In all conditions the heavy B-pool disappeared within 2.5 h of chase (Figure 3D, Figure S4D), being degraded or matured into the C-pool. Similar to previous studies [7, 11, 29], only a fraction (∼19 %) of the initial heavy B-pool of WT-CFTR was converted to a heavy C-pool (2.5 h chase), while <1 % of the heavy B-pool of F508del-CFTR was converted at this time point (Figure 3D-E). TEZ or ELX correction of F508del-CFTR increased B-to-C conversion rates to 2.4 % or 6.7 %, respectively, while IDOR-3 or IDOR-4 resulted in conversion rates of 24-31%, which were thus higher than observed for WT-CFTR. We conclude that type IV correctors elevate F508del-CFTR ER folding efficiency to or even beyond WT levels, i.e., fully compensate the F508del-induced folding defect.

To determine how correctors affected stability of already folded F508del-CFTR, we labeled the whole CFTR pool with light amino acids and performed a chase with heavy amino acids. Correctors at their maximally effective concentrations were either present during both pulse and chase periods or washed out after the pulse (Figure 3F). For WT-CFTR the light C-pool decayed slowly (t_1/2_=18 h), whereas the light C-pools of F508del-CFTR corrected with TEZ, ELX, IDOR-3 or IDOR-4 decayed faster (t_1/2_ of ∼5-6 h), irrespective of corrector type and presence, suggesting that none of the correctors were efficient CFTR stabilizers (Figure 3G, Figure S4E, upper panels). Wash-out of the correctors (except IDOR-4, still showing 50% of heavy C-pool) was confirmed by the lack of new heavy C-pool formation in the washed conditions (Figure S4E, lower panels). Similar results were obtained in reconstituted bronchial epithelium (Figure S4F-G) where the WT-CFTR C-pool displayed a t_1/2_ of ∼15 h while the IDOR-3- and IDOR-4-corrected F508del-CFTR C-pool had a t_1/2_ of ∼5-7 h. For TEZ- and ELX-treated samples, t_1/2_ values could not be determined as the signal was too low. In contrast to previous techniques, which lack wash-out controls and use cycloheximide to block protein neosynthesis [7, 11], this highly sensitive pulse-chase approach showed that neither type-III, -I or the new type-IV correctors alter F508del-CFTR post-folding stability. However, since type-IV correctors promote F508del-CFTR ER folding to up to 170 % of wildtype efficiency, a steady state F508del-CFTR expression of almost wildtype levels is reached (Figure 1F).

### Type-IV correctors overcome thermal instability during F508del-CFTR synthesis and require MSD1-NBD1-R-MSD2 as minimal substrate

We further dissected type-IV correction mechanistically. Rescue of ER folding efficiency to WT levels can be achieved by simultaneous correction of the thermal and interface instability in F508del-CFTR, e.g., by introducing suppressor mutations that thermally stabilize NBD1 (i.e. G550E.R553M.R555K or “3TS”) in combination with mutations that increase NBD1-MSD2 interface interactions (R1070W or “RW”) [8, 31–33]. Indeed, HEK-F508del-CFTR.3TS cells expressed 6-fold more C-band CFTR than HEK-F508del-CFTR cells and the addition of the RW mutation yielded a further 2-fold increase for an overall 12-fold increase (Figure 4A-B). We applied the correctors at maximally effective concentrations as assessed for this specific cell type and serum concentration (Figure S5A-B). In F508del-CFTR cells, IDOR-3/IDOR-4 induced 9-/12-fold increases in C-band expression (Figure 4A-B), i.e., IDOR-4 had the same efficiency as the combined introduction of 3TS and RW mutations. Combining IDOR-3 or IDOR-4 with 3TS mutations only marginally improved C-band expression (∼1.5-fold) and the introduction of the RW mutation had no additional effect. In contrast, TEZ and ELX, strongly increased C-band expression (∼4-fold) after introduction of 3TS mutations, indicating that TEZ and ELX do not rescue thermal instability (Figure 4 A-B) [8, 34]. Taken together, type-IV correctors have the ability to fully overcome the F508del-CFTR folding defects yielding efficiencies comparable to the suppressor mutations that remedy both thermal and interface instability during folding.

**Fig. 4:**
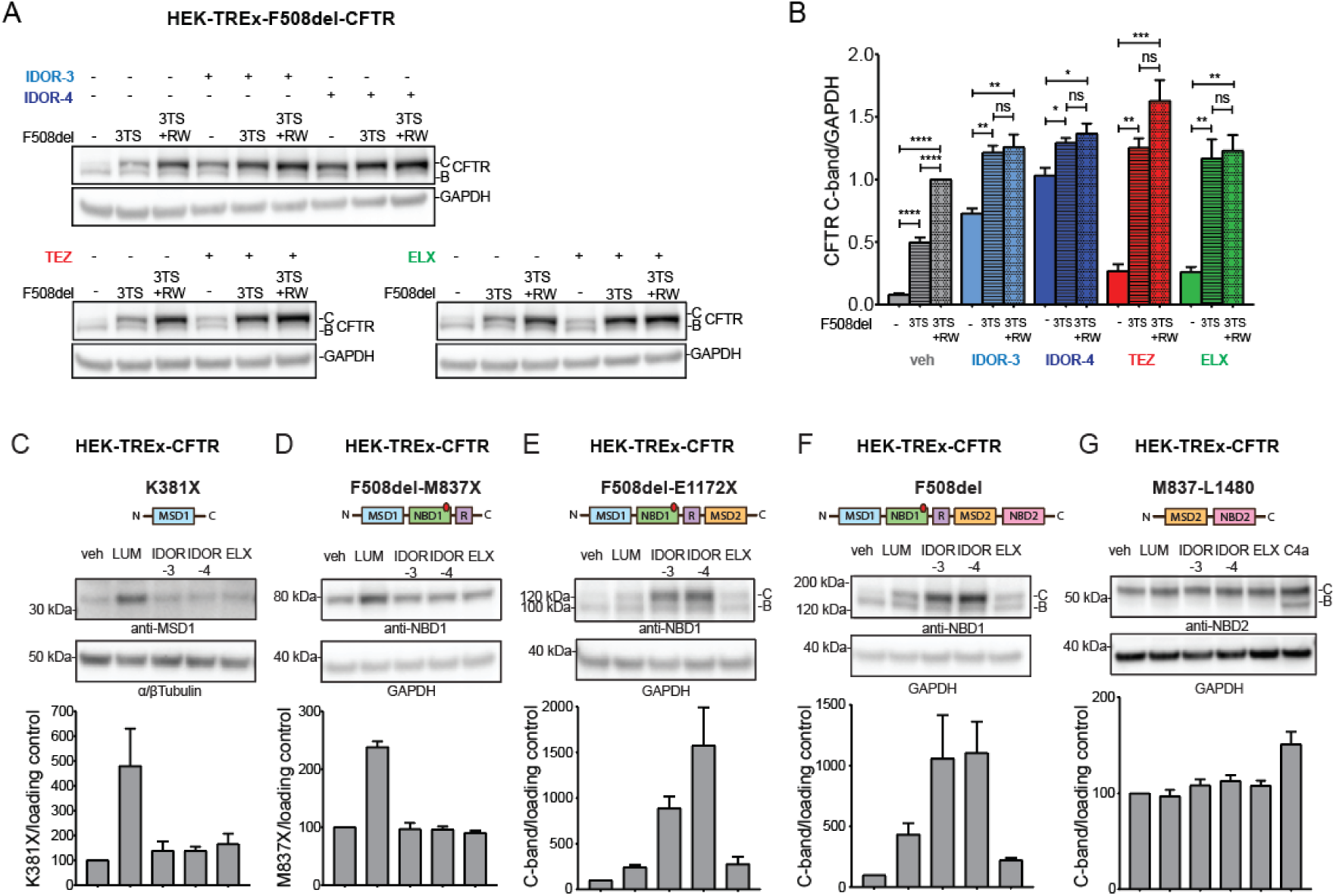
Type-IV correctors restore native folding by overcoming thermal instability during F508del-CFTR synthesis and require MSD1-NBD1-R-MSD2 as minimal substrate. (**A**) CFTR correction (immunoblotting) in HEK-TREx cells expressing either F508del-CFTR, or thermostabilized F508del-CFTR-G550E.R553M.R555K (3TS), or thermostabilized and interface-stabilized F508del-CFTR-G550E.R553M.R555K.R1070W (3TS+RW). Cells were treated for 48 h with DMSO or maximally effective concentrations of either IDOR-3 (2 µM), IDOR-4 (0.4 µM), TEZ (2 µM), or ELX (0.4 µM). Representative images of 3 experiments. **(B)** CFTR C-band intensities in (A) normalized for GAPDH and further normalized to the C-band intensity of vehicle-treated F508del.3TS.RW cells (n=3). **(C-G)** Levels of truncated CFTR constructs (immunoblotting) expressed in HEK-TREx cells after 40 h of treatment with vehicle or maximally effective concentrations of either LUM (2 µM), IDOR-3 (2 µM), IDOR-4 (0.4 µM), ELX (0.4 µM), or Corrector4a (10 µM). Detection antibodies target either MSD1 (C), NBD1 (D-F), or NBD2 (G). Intensities of CFTR fragment bands are shown in the graphs (C-band for E-F-G), normalized for the loading controls (n=3 for C-F, n=4 for G). Data in (B), (C), (D), (E), (F), (G) are means ± SEM of the indicated number of independent experiments. In (B), a one-way ANOVA test was performed between the three different mutants within every compound treatment group; ns: non-significant: *: p-value < 0.05; **: p-value < 0.01; ***: p-value < 0.001; ****: p-value < 0.0001. See also Figure S5.

To pinpoint the CFTR protein domains involved in type-IV correction, we generated HEK cells expressing CFTR fragments of increasing size (Figure 4C-G) as previously described [35]. Expression of MSD1 (K381X-CFTR, Figure 4C) increased 5-fold after LUM treatment in agreement with the recently published binding site and mode of action of type-I correctors, which bind to and increase the stability of MSD1 [12–15]. IDOR-3, IDOR-4, and ELX had no effect. Similar results were obtained for the MSD1-NBD1-R fragment (F508del-M837X-CFTR, Figure 4D). A clear change was observed for the MSD1-NBD1-R-MSD2 fragment (F508del-E1172X-CFTR, Figure 4E), which was corrected and glycosylated to produce a C-band by treatment with LUM or ELX (2-3-fold over baseline) and especially IDOR-3 or IDOR-4 (10-15-fold over baseline). Similar effects were observed for full-length F508del-CFTR (Figure 4F). None of these correctors affected expression of the MSD2-NBD2 fragment (M837-L1480-CFTR, Figure 4G) while type-II corrector C4a, proposed to promote NBD2 folding [36], increased the expression of its glycosylated and non-glycosylated forms. These data confirm that type-I correctors act on MSD1 and type-II correctors act on NBD2. Importantly, they demonstrate that type-IV and type-III correctors require an almost full length CFTR substrate including MSD2, likely because they promote the challenging assembly of MSD2 with the preceding domains.

### Type-IV correctors bind to the CFTR lasso domain

To covalently label the type-IV corrector binding site, three photo-activatable aryl-azide macrocycle probes (IDOR-5A, -6A, -6B) were developed from non-azide parent molecules IDOR-5 and IDOR-6 (Figure 5A). These 5 compounds were designed with ease-of-synthesis considerations and showed pharmaco-chaperone activity comparable to IDOR-3 and IDOR-4 (Figure S6A-C). In buffer containing primary amines (Tris), the azide-containing probes were completely turned over within 5 min of UV illumination demonstrating their UV-inducible reactivity (Figure S6D). U2OS-F508del-CFTR cells treated with the three photo-activatable probes or the two non-azide controls were exposed to UV illumination or not. Proteins were separated by SDS-PAGE, the CFTR-containing gel regions were excised and trypsin-digested for LC/MS analysis (Figure 5A). In four independent experiments an average of 73 CFTR peptides were detected corresponding to 53% of the protein (Figure S6E-F). Only 2 out of these 73 peptides were significantly (with multiple testing correction) reduced under UV exposure when comparing each photoactive probe with its corresponding non-active parent macrocycle: compared to parent IDOR-6, IDOR-6A reduced by ∼50% the abundance of the overlapping peptides _15_LFFSWTR_21_ (p.adj=0.0002) and _15_LFFSWTRPILR_25_ (p.adj=0.00001) (Figure 5B, S6G). No other peptide showed significant down-regulation (Figure S6G-H). The results for these two peptides without multiple testing correction confirmed significant reductions for IDOR-6A versus IDOR-6 (with UV) and, in addition, for IDOR-6A with UV versus without UV, and showed no differences for the other two aryl-azides IDOR-6B and IDOR-7A (Figure 5B). Thus, cross-linking of IDOR-6A occurred between amino acids (aa) 14 and 21 (K14 is recognized by trypsin), i.e., at the N-terminus of CFTR, within lasso helix-1 (Lh1) (Figure 5C). Interestingly, neither type-I, -III or -IV correctors could rescue F508del-CFTR in which aa14-21 had been deleted (Figure S7A), suggesting that aa14-21 are crucial for the CFTR folding process. In contrast, deletion of aa14-21 in the isolated MSD1 domain led to an increased MSD1 expression (Figure S7B) suggesting a destabilizing effect of aa14-21 on the isolated domain itself. These data indicate that type-IV correctors interact with a sequence in Lh1 that plays an essential role in folding full-length CFTR.

**Fig. 5.**
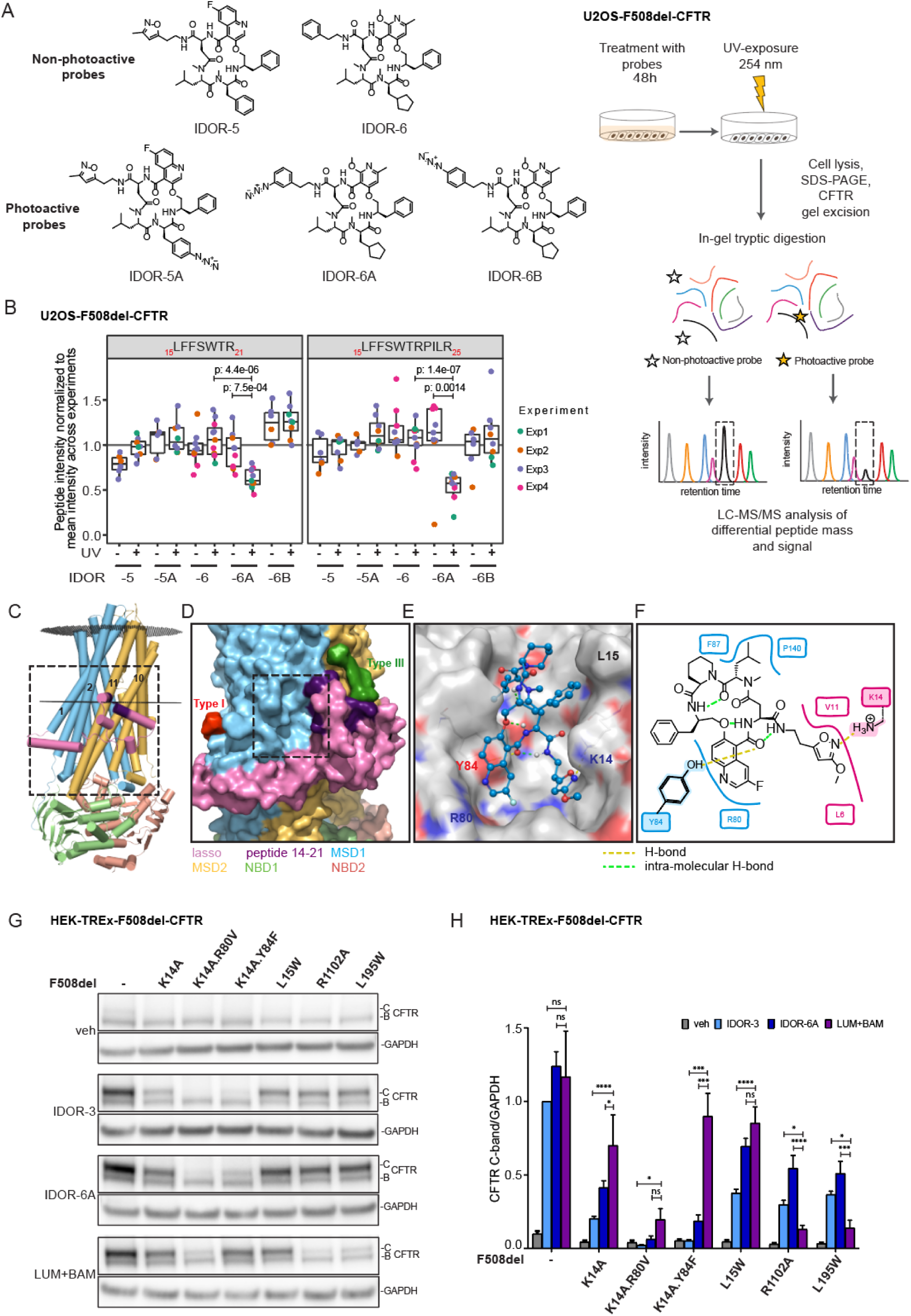
Type-IV correctors bind to the CFTR lasso domain. **(A)** Determining the type-IV corrector binding site on CFTR: structures of the photoactive probes and scheme of the UV-crosslinking experiments with subsequent CFTR peptide mapping using U2OS-F508del-CFTR cells. **(B)** Abundance of two specific CFTR peptides _15_LFFSWTR_21_ and _15_LFFSWTRPILR_25_ from cells treated for 48 h with 2 µM of the non-photoactive probes IDOR-5 or IDOR-6 or the corresponding photoactive probes IDOR-5A, IDOR-6A or IDOR-6B, and then exposed or not to UV light for 5 min. The signal for each peptide was normalized to the amount of CFTR in that sample using the average response factor of that peptide across all experiments. The scatter plot represents 4 independent experiments with dots of the same color representing technical replica. Significant p-value after one-sided t-test analysis (not adjusted for multiple testing) was achieved only for IDOR-6A versus IDOR-6 (both with UV) and IDOR-6A with UV versus without UV. **(C)** Cylinder view of the F508del-CFTR channel in the phosphorylated open state (PDB:8EIQ). The plasma membrane border is depicted with dark grey discs. The crosslinked lasso domain peptide (including K14) _14_KLFFSWTR_21_ is depicted in purple. TM1, 2, 10, and 11 are highlighted. **(D)** Surface view of the cavity in the vicinity of the crosslinked peptide which faces TM1/2 and can accommodate active macrocycles. Type-I and type-III correctors in their published binding poses [21] are depicted in red and green, respectively. **(E-F)** 3-dimensional (E) and 2-dimensional (F) view of IDOR-3 bound in the CFTR cavity as determined by docking studies. Polar amino acid side chains in (E) are indicated in blue, red and yellow. H-bonds between IDOR-3 and CFTR are indicated as yellow dashed lines. IDOR-3 intramolecular H-bonds are depicted as green dashed lines. In (E), Y84 and K14 side chains are highlighted. **(G)** Immunoblot analysis of corrector activities after site-directed mutagenesis of key amino acids in the proposed type-IV corrector binding site: F508del-CFTR constructs containing one or two additional mutations in the site (K14A, K14A.R80V, K14A.Y84F or L15W) or the ELX binding site (R1102A) or the LUM binding site (L195W) were expressed in HEK-TREx cells, then treated for 40 h with 2 µM of each indicated corrector. Representative images from at least 4 independent experiments per mutant and per corrector treatment. **(H)** Quantification of F508del-CFTR C-band intensities in (F) normalized for GAPDH (n=4-10). Data in (G) are means ± SEM of the indicated number of independent experiments. One-way ANOVA test between the different compound treatments within every mutant, the comparison between [LUM+BAM] and each of the two macrocycles is shown: ns: non-significant: *: p-value < 0.05; ***: p-value < 0.001; ****: p-value < 0.0001. See also Figure S6 and Figure S7.

Considering the macrocycle size, two binding sites adjacent to peptide14-21 were conceivable (Figure 5C-D). One site faces MSD2-TM11 (Figure 5D), and is identical to the recently described binding site for type-III correctors, here shown with ELX bound [21]. The remaining other site faces MSD1-TM1/2 (Figure 5D, box) and is a cavity at the membrane-cytosol interface that can accommodate active macrocycles such as IDOR-3 with high shape complementarity and involving polar interactions with at least two polar amino acids (K14 in Lh1, Y84 in MSD1) as determined by molecular docking (Figure 5E). This is exemplified for IDOR-3, in which the isoxazole moiety forms an H-bond with the side chain amine of K14, and one central-ring carbonyl forms an H-bond with the hydroxy group of the Y84 side chain. The other interactions are lipophilic and based on optimized shape complementarity. Thus, IDOR-3 is proposed to act as a bridge between MSD1-TM1/2 and Lh1. Interestingly, all three non-methylated macrocyclic amides are involved in intramolecular H-bonds, placing the macrocycle in a favorable, low energy conformation (Figure 5E-F). This binding mode hypothesis was transferable to the other active macrocycles and used to rationalize why IDOR-6A was the only aryl-azide that could be cross-linked to CFTR: In IDOR-5A, the aryl-nitrene (activated intermediate) is too distant from the CFTR surface (Figure S7C). In IDOR-6A, the aryl-*meta*-nitrene is optimally placed to covalently bind to the K14 amine (Figure S7E-F), while the aryl-*para*-nitrene in IDOR-6B is not (Figure S7D).

Next, this site was explored by introducing single or double mutations to the 3 polar amino acids K14, Y84 and R80 within F508del-CFTR to either disrupt polar interactions (K14A, K14A.R80V, K14A.Y84F) or occlude part of the site (L15W). Known mutations abolishing type-III (R1102A) and type-I (L195W) correction were introduced as controls [12, 21]. Cells were then treated overnight with different corrector types and combinations at their maximally effective concentrations as assessed for this specific cell type and serum concentration (Figure S5A-B). Without correctors, CFTR mutants were expressed only as B-bands (Figure 5G-H). The double and triple mutants generally displayed slightly lower correctability as compared to F508del-CFTR (Figure S7G-H). Our analysis of the different mutations deliberately focused on comparing the activity of macrocycles (IDOR-3, IDOR-6A) versus [LUM+BAM = type-I + type-III correction], because [LUM+BAM] and macrocycles reached similar correction of F508del-CFTR in this cell type (Figure 5F-G) allowing for clear interpretation of mutation effects: F508del.L195W-CFTR and F508del.R1102A-CFTR were only weakly corrected by [LUM+BAM] compared to the macrocycles (∼3- to 5-fold lower), as expected for mutations disrupting type-I or -III corrector binding. F508del.L15W-CFTR showed little difference in correction by macrocycles versus [LUM+BAM]. In contrast, F508del.K14A-CFTR displayed a weaker correction by macrocycles (2-4-fold), as compared to [LUM+BAM]. A striking selective 5-10-fold loss of correction for macrocycles versus [LUM+BAM] was observed in the F508del.K14A.Y84F triple mutant. The F508del.K14A.R80V triple mutant was barely correctable by any corrector, likely due to loss of an essential intra-protein salt bridge involving R80. These data (Figure 5G-H) and further data in the supplement (Figure S7G-H) strongly suggest that type-IV correctors bind in the cavity between Lh1 and MSD1-TM1/2, interacting with the polar amino acids K14 and Y84.

### Type-IV correctors address non***-***F508del CFTR folding mutations

The observed site of action of type-IV correctors which is distant from the actual F508 deletion suggested that other CFTR folding mutations might be rescued. We expressed the ten most frequent CF-causing CFTR folding mutations (https://cftr2.org) (Figure 6A-B). Using maximally effective concentrations (Figure S5C-D) we tested IDOR-3, IDOR-4 and [TEZ+ELX] (stand-alone TEZ or ELX treatment showed no/little correction in many mutants, Figure S5C-D). [TEZ+ELX] were highly effective in L206W, R347P, S945L, moderately effective in M1101K and R1066C and not effective in G85E, I507del, R560T, V520F and N1303K (Figure 6C-D, Figure S5C-D). In contrast, macrocycles were highly effective in all mutants except N1303K where only minor activity was detectable (Figure 6C-D) showing that the allosteric mechanisms promoting MSD1/2-NBD1 folding only weakly extend to the C-terminal NBD2. Taken together, type-IV correction can address many non-F508del-CFTR folding mutations including the currently not treatable I507del, R560T, V520F and R1066C, demonstrating an unprecedented potential for this novel corrector mechanism.

**Fig. 6.**
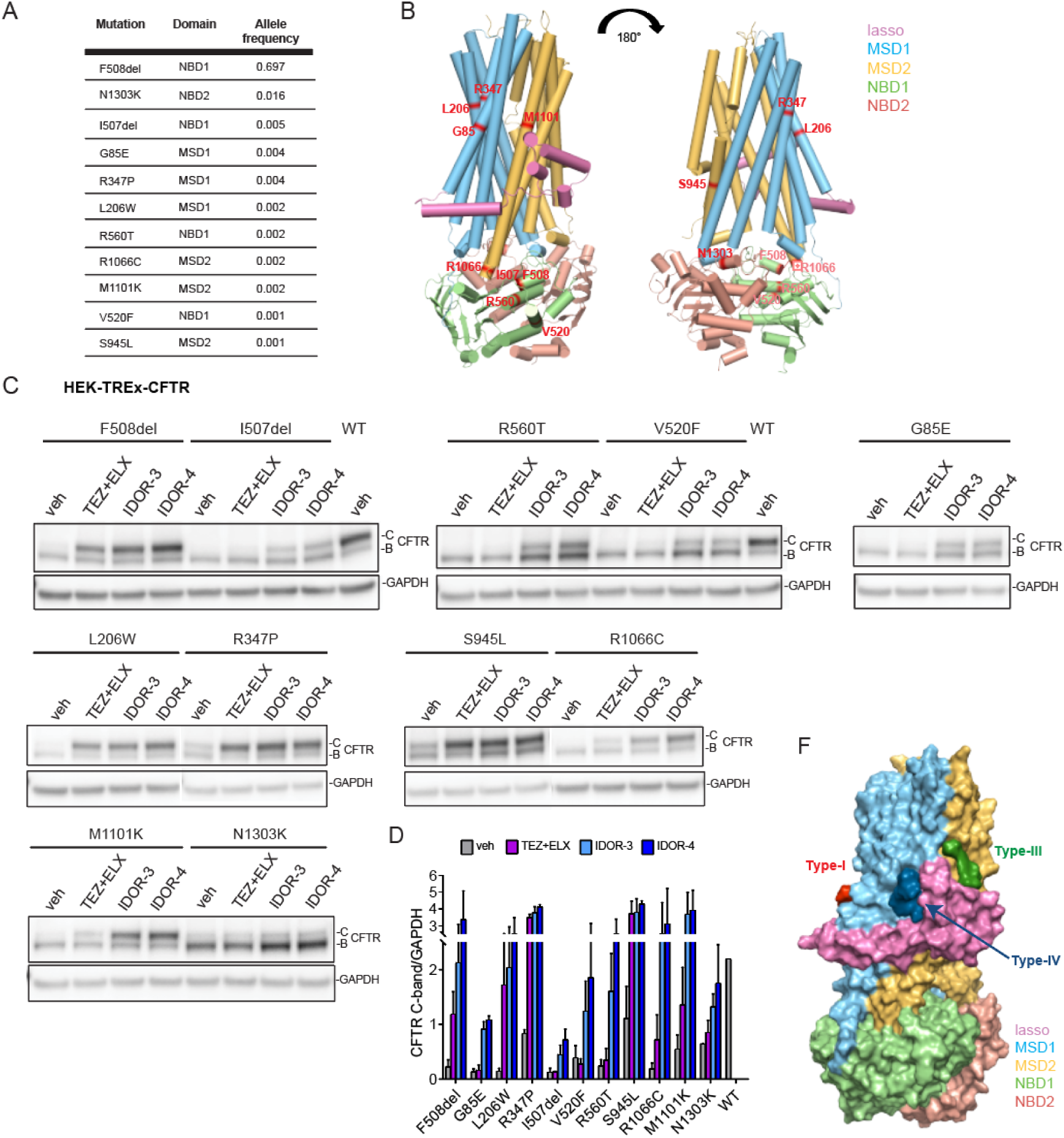
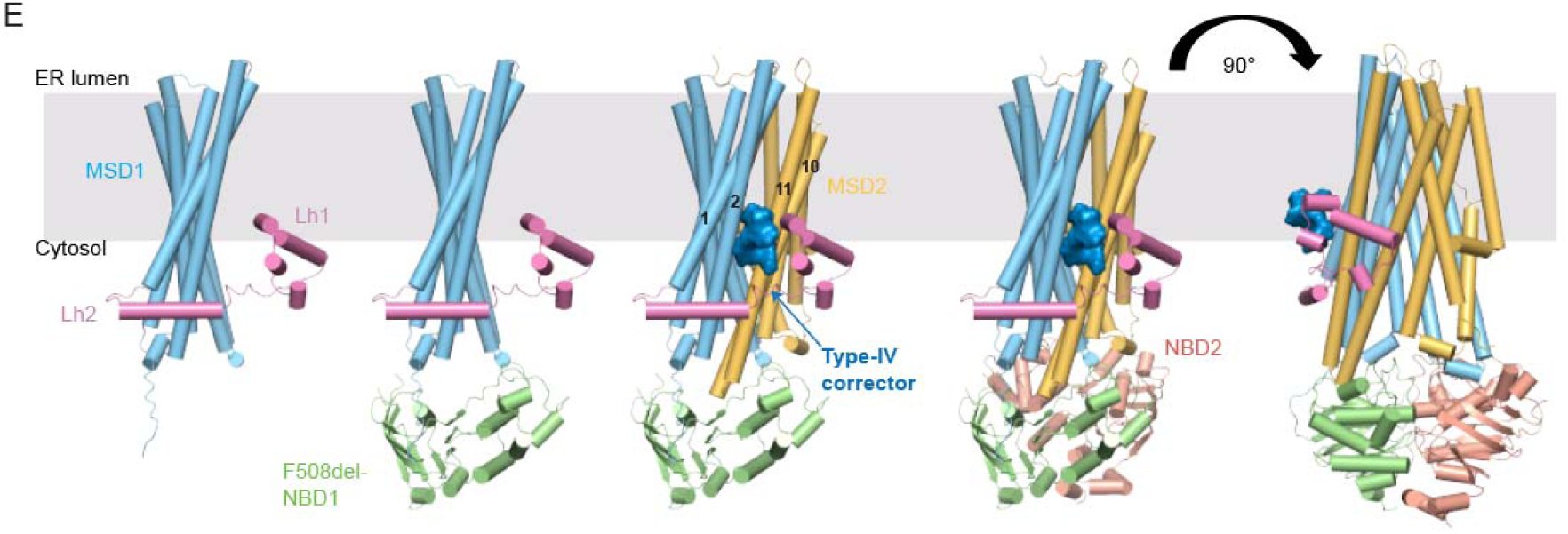
Type-IV correctors address non-F508del CFTR folding mutations. **(A)** Most frequent class-II folding mutations, their domain localization and allele frequency among the CF patient population. **(B)** Cylinder view of the WT-CFTR channel in the phosphorylated-open state (PDB:6MSM) with the amino acids of the corresponding mutations colored in red. **(C)** CFTR correction (immunoblotting) in HEK-TREx cells expressing 11 different class-II folding mutations (see A, B) and treated for 48 h with DMSO or maximally effective concentrations of IDOR-3 (2 µM) or IDOR-4 (0.4 µM) or a combination of TEZ (2 µM) + ELX (0.4 µM). Representative images from 2 experiments. **(D)** CFTR C-band intensities in (C) normalized for GAPDH (n=2). Data represent mean ± SEM of two independent experiments. **(E)** Model depicting type-IV corrector binding to the Lh1-MSD1-MSD2 interface to support the challenging interdigitation of transmembrane helices which occurs late in translation. **(F)** Model of simultaneous type-I, -III and -IV corrector binding to CFTR. See also Figure S5.

## DISCUSSION

CFTR modulators have significantly improved life quality and expectancy for many CF patients. Since single-corrector treatments showed limited effects the current standard of care for F508del-CFTR patients combines type-I corrector TEZ and type-III corrector ELX to mediate - together with the potentiator IVA – a substantial improvement of CFTR function. Current consensus is that CFTR correctors must act in concert to achieve clinically effective correction, however CFTR folding and trafficking is still not recovered by more than ∼50%. This deficit warrants the search for correctors with novel, complementary mechanisms, to be used in addition to existing corrector types or even as highly effective stand-alone treatments. Compared to type-I, -II, and -III correctors, the macrocyclic type-IV correctors from our drug discovery program display unprecedented efficacy in promoting folding and trafficking of F508del-CFTR in cell lines and reconstituted patient tissue. In addition to having therapeutical potential to address the F508del folding mutation as well as many others, type-IV correction also sheds more light onto the CFTR folding process because the interdomain macrocycle binding site pinpoints a CFTR region highly relevant to its folding.

Three complementary approaches using photo-activatable probes, molecular modeling/docking and site-directed mutagenesis resulted in the identification of a highly probable binding site of type-IV correctors. Crosslinking yielded only a single CFTR peptide pair that was consistently reduced under UV exposure by only one (the aryl-*meta*-azide) of the three cross-linkable probes and suggests a highly specific macrocycle interaction with CFTR peptide14-21. The N-terminus of CFTR (aa1-80) is highly conserved among species and ABC-C transporters [37, 38] and is an important contributor to folding. Lasso helix-2 (Lh2, aa46–61) interacts with MSD1-TM1-4 and the N-terminus of NBD1 to create a first important domain assembly unit [13]. The function of lasso helix-1 (Lh1, aa11-29) remained largely unknown until the recent disclosure of the cryo-EM structure of F508del-CFTR bound to ELX, TEZ and IVA in which ELX is positioned in a void between Lh1 and MSD2-TM11 with Lh1 packing against MSD1-TM1/2 and MSD2-TM10/11 [21, 38]. Our study highlights a central role of Lh1 in promoting folding of mutated and probably also WT-CFTR: the deletion of aa14-21 abolishes the folding capacity of F508del-CFTR in presence of any corrector (Figure S7A) while the same deletion in the isolated MSD1 strongly increases domain expression (Figure S7B). This contrasting behavior suggests a central chaperoning role of Lh1 in MSD1-MSD2 interdomain assembly likely assisting in the challenging interdigitation of freshly synthesized TM10/11 of MSD2 into the TM1/2/3/6 helix bundle of MSD1. Accordingly, type-IV correctors that interact with Lh1, exert their stabilizing effects only if the CFTR sequence spans the region from the N-terminus to MSD2 (Figure 4C-G), in contrast to type-I correctors that promote MSD1 intra-domain stability [12–15, 39] (Figure 4C). Thus, it is likely that macrocycles, by binding in the cavity between Lh1 and MSD1-TM1/2, stabilize a late translational intermediate complex of Lh1, MSD1 and MSD2 (Figure 6E). Type-III correctors, according to [21] and our data have a related but not identical mode of action (hence the additivity) by stabilizing Lh1 from the other side which faces MSD2-TM11, but with lower efficacy. We conclude that Lh1-mediated interdomain assembly of MSD1 and MSD2 is crucial for CFTR folding, and pharmaco-chaperones acting on this step -the type-III and the more efficient type-IV correctors - can allosterically overcome the need of a direct rescue of NBD1 instability during folding (Figure 4A-B). The non-overlapping nature of corrector type-I, -III and -IV binding sites (Figure 6F) plausibly explains their additive behavior in correcting CFTR folding. Taken together, characterizing type IV correction has uncovered a short N-terminal amino acid stretch – Lh1 - with a central role in CFTR inter-domain assembly. Using this highly effective allosteric mechanism, type-IV correctors can address many severe folding mutations in MSD1, NBD1 and MSD2 that are not correctable with other corrector types (Figure 6A-D), and even slightly promote folding of WT-CFTR.

Steady state levels of cell surface CFTR are governed by CFTR folding/trafficking efficiency (ER) and CFTR post-folding half-life (PM). Combining the SILAC pulse-chase technique with SDS-PAGE, B/C-band excision and CFTR peptide LC/MS analysis, allowed us to precisely quantify these processes and their modulation in CFBE cells. WT-CFTR folding efficiency (∼20%) and protein half-life (t_1/2_=18 h) were in agreement with previous reports as were those of F508del-CFTR (<1% folding efficiency) (Figure 3D-E). Folding efficiency of the analyzed correctors was between 2% and 31% (Figure 3D-E) displaying the same rank order as determined in Western blotting of F508del-CFTR steady state levels (Figure 1E-F). This indicates that the difference in corrector efficacy on steady-state CFTR levels is a consequence of their different CFTR ER folding efficacies reaching more than wildtype levels with the best type-IV correctors (31%). However, no corrector (including type-IV correctors) prolonged the stability of already folded F508del-CFTR (t_1/2_∼5 h; Figure 3G) as determined with our novel pulse-chase technique avoiding cycloheximide and including corrector washout controls. Thus, correctors promote the stability of CFTR folding intermediates during translation, but do not address the stability deficit of mature F508del-CFTR. Still, steady state expression of macrocycle-corrected F508del-CFTR almost reaches WT levels, likely because ER folding efficiencies are increased considerably beyond wildtype levels.

In conclusion, type-IV correction optimally supports an essential intramolecular folding mechanism involving the previously underestimated lasso helix-1, and thus fully normalizes F508del-CFTR folding and trafficking to essentially wildtype levels, reaching efficacies that are superior to the previously known type-I, -II and -III corrector mechanisms. Since this unprecedented efficacy is recapitulated in reconstituted CF patient tissue, type-IV correctors whose drug-like properties will be described in due course, could represent a promising new treatment modality for CF patients with CFTR folding mutations.

## MATERIALS AND METHODS

### Compounds

Lumacaftor was purchased from Astatech, Tezacaftor from Selleckchem, Ivacaftor from Combi-Blocks and Corrector 4a from Manchester Organics. All other compounds used in this study were synthetized at Idorsia Pharmaceuticals, Ltd (Allschwil, Switzerland). The synthesis of correctors IDOR-1 to IDOR-4 is described in patent WO2022194399A1.

### Cell line origin and culture

PathHunter® U2OS-F508del-CFTR MEM-EA and PathHunter® U2OS-ADRB2(W158A) ENDO-EA cells (abbreviated U2OS F508del-CFTR and U2OS ADRB2[W158A]) were obtained from DiscoverX (catalog 93-0987C3 and 93-0986C3, respectively) and were propagated in their culture medium (AssayComplete Cell Culture Kit 103, DiscoverX) according to manufacturer instructions.

Cystic Fibrosis Bronchial Epithelial (CFBE41o-) cells stably expressing the human wild-type (WT) CFTR or F508del-CFTR mutant (generated in [28], were a kind gift from Dr. J.P.Clancy (University of Alabama, Birmingham) and cultured on fibronectin-coated plates (FNC-coating mix; Enzo) in Minimum Essential Medium (MEM) with Earl’s Salt (ThermoFisher Scientific) containing 10% of heat inactivated fetal bovine serum (FBS; Gibco), 100 U/ml penicillin, 100 mg/ml streptomycin (ThermoFisher Scientific) and 2µg/ml puromycin (Sigma-Aldrich).

The human CFTR expressing HEK cell pools with the mutations described below, were generated by integrase-mediated homologous recombination of the CFTR sequence into the T-REx HEK293 background (ThermoFisher Scientific). This expression system allows the tetracycline-inducible expression of genes of interest from one defined insertion site. Parental T-REx-HEK293 cells were cultivated in growth medium: Dulbecco’s modified Eagle’s medium (DMEM) + GlutaMAX-I (#31966, Life Technologies), 10% Dialyzed FBS (Gibco, #26400), 100 U/ml penicillin, 100 *µ*g/ml streptomycin, 0.1 mg/ml hygromycin B (Life Technologies), 5 μg/ml blasticidin (Invitrogen, #R210-01), and 0.1 mM nonessential amino acid solution (NEAA, Invitrogen, #11140-035). Recombinant T-REx-HEK-CFTR cells were generated by lipofectamine transfection (Life Technologies) of plasmids containing mutated versions of CFTR (RefSeq NM_000492), and then cultivated in selection medium (growth medium containing 1 mg/ml Geneticin [Invitrogen, #10131-027] instead of hygromycin). The plasmid constructs were synthetized by GENEART (ThermoFisher Scientific) and designed with the wildtype CFTR intron 5-6 (880nt) to prevent leaky expression issues during the cloning process in bacteria. For evaluation of corrector efficacies on second-site suppressor mutants, the following mutations were introduced into the CFTR.F508del gene sequence: G550E.R553M.R555K and G550E.R553M.R555K.R1070W. The following plasmids expressing different truncated versions of CFTR.F508del gene were generated: K381X, delF508-M837X, delF508-E1172X and M837-L1480. For the evaluation of different binding sites by site-directed mutagenesis, the following mutations were introduced individually into the CFTR.F508del gene: del14-21, K14A, K14A.R80V, K14A.Y84F, L15W, R1102A, L195W. Deletion of the peptide stretching amino acids 14-21 was also introduced into truncated K381X: del14-21.K381X. To assess corrector capacity to stabilize different CF-causing, class-II folding mutants other than F508del, the following mutations were introduced to the CFTR wildtype gene sequence: G85E, L206W, I507del, V520F, R560T, S945L, R1066C, M1101K, N1303K and R347P. For co-potentiation experiments in the YFP-quenching assay, the CF-causing, class-III gating mutation G551D was introduced to the CFTR wild-type gene sequence. For the halide-sensitive YFP quenching assay, U2OS F508del-CFTR or TREx-HEK-G551D-CFTR cells were transduced with lentiviral particles rLV.EF1. F46L/H148Q/I152L-YFP (Vectalys, FlashTherapeutics) carrying a mutated halide-sensitive version of YFP [24]. Briefly a suspension of 5.000.000 cells was added to a mix of lentiviral vectors (final concentration 2.7x10^9^ TU/mL) mixed with 4µg of Polybrene (Sigma-Aldrich). Then cells were seeded and expanded in their respective growth medium.

### Pharmacochaperone trafficking assay

This assay was adapted from the DiscoverX PathHunter® pharmacochaperone trafficking assay which is based on the enzyme fragment complementation technique. In brief, the U2OS-F508del-CFTR cell line (DiscoveRx, #93-0987C3) is engineered to co-express (i) human F508del-CFTR tagged with a Prolink (PK =short β-galactosidase fragment) and (ii) the remainder of the β -galactosidase enzyme (Enzyme Acceptor; EA) localized to the plasma membrane. Incubation with compounds that increase trafficking of PK-tagged F508del-CFTR to the plasma membrane will lead to complementation of the EA fragment to form a functional ß-galactosidase enzyme which is quantified by a chemiluminescence reaction reporting cell surface CFTR expression. The same principle applies to the counter-screen using U2OS-ADRB2(W158A) cells (DiscoveRx, #93-0986C3). Freshly thawed U2OS-F508del-CFTR and U2OS-ADRB2(W158A) were seeded at 3500 or 5000 cells/well respectively in 20 µl/well of cell assay medium (McCoy [Life Technologies], 10 % FBS, 100 U/ml penicillin, 100 mg/ml streptomycin) in a 384-white-low volume plate (Corning). Pre-dilutions of compounds in DMSO were prepared in a 384-well polypropylene plate using an automated liquid-handling system. Dilution series were then transferred into cell assay medium in a polypropylene 384-well plate (Greiner) to obtain 5x concentrated aqueous working stocks. Five hours after cell seeding, 5 µl/well of 5x working stocks were added to the 20 µl/well of cells for an incubation over-night at 37°C. The next day, after 2 h at room temperature, the cell plates were incubated with 10µl of Flash detection reagent (#93-0247, DiscoverX) for 30 min and chemiluminescence signals were detected with a microplate reader (Synergy MX, Agilent BioTek). Chemiluminescence data were normalized to the signal of basal vehicle-treated cells and mean chemiluminescence values were converted to concentration-response curves for determination of EC_50_ and E_max_ values (GraphPad Prism).

### Reconstituted primary human bronchial epithelial tissues

Reconstituted tissues from CF and non-CF donors were either directly purchased as fully differentiated air-liquid interface culture from Epithelix SA (Geneva) for direct experimental use or generated in our laboratories from passage-1 human airway epithelial cells (hAEC; obtained from Epithelix SA) using the differentiation protocols and reagents provided by Stemcell Technologies (PneumaCult^TM^ reagent series). In-house generated cultures were derived from three homozygous F508del-CFTR patients: CF patient#1 (EP57AB-CFAB060901, Epithelix, female, 21 years-old), CF patient#2 (EP57AB02-CFAB0452, Epithelix, male, 32 years-old), CF patient# 3 (#28388-0000450918, Lonza, male, 25 years-old) and four non-CF donors with no pathology reported: non-CF#1 (EP51AB-02AB079301, Epithelix, male, 62 years-old), non-CF#2 (EP51AB-02AB0834.01 Epithelix, male, 71 years-old), non-CF#3, (EP51AB-02AB0839.01, Epithelix, male, 59 years-old), non-CF#4 (CC-2540S -20TL356517, Lonza, female, 48 years-old). hAEC cells were cultured for several days until reaching 90-95% confluency in complete PneumaCultExPlus medium. Cells were then washed with PBS and dissociated with ACF enzymatic dissociation solution, followed by addition of ACF enzyme inhibition solution. Cells were seeded onto single transwell inserts (polyester membranes, Corning, #CLS3470 in 24-well plates, #Corning 3524) sitting in 600 µl/well of complete PneumaCultExPlus medium at a density between 30.000-50.000 cells/insert and cultivated submerged for several days. Once cells reached 90-95% confluency, medium was removed from the apical side to expose cells to air, while the medium on the basal side was replaced with 600 µl/well PneumaCult-Air-Liquid-Interface (ALI) maintenance medium. Basal medium was exchanged every three days for four weeks till full differentiation was reached as indicated by ciliary beating, mucus production and tight junction formation (ZO-1 staining).

### Treatments with correctors for immunoblotting

For testing concentration-response effects of correctors on rescuing F508del-CFTR C-band expression in U2OS-F508del-CFTR cells, cells were treated as described for the pharmacochaperone trafficking assay. CFBE-F508del-CFTR or CFBE-WT-CFTR were seeded at 140.000 cells/well in a FNC-coated 24-well plate for a day and then incubated for 24 h with ascending concentrations of correctors diluted in growth medium deprived of FBS but supplemented with 1% Human Serum Albumin (HSA, Sigma-Aldrich, #A1653). The same protocol was used for treatment with proteasome inhibitor MG-132 (Sigma-Aldrich). For CFTR expression kinetics and reversibility upon wash-out, CFBE-F508del-CFTR cells were seeded and treated as above with fixed concentrations of correctors for 2, 4, 8, 24, 27, 31 and 48 h. At the 24 h time point, cells for wash-out analysis were washed twice for 10 min with CFBE growth medium and then incubated for 3 h, 7 h or 24 h in 1 % HSA-medium without correctors. For all experiments with primary bronchial ALI tissues, correctors were added to the basal medium for 24h. For all experiments with isogenic T-REx-HEK cells expressing various CFTR mutants, cells were seeded at 200.000 cells/well in a 24-well plate for a day and then incubated for 40 h with ascending or fixed concentrations of correctors diluted in growth medium supplemented with 10 ng/ml tetracycline (tetracycline hydrochloride, Sigma, #T7660). For T-REx-HEK cells expressing truncated versions of CFTR, 100 ng/ml of tetracycline was used to ensure sufficient expression levels.

### Immunoblotting

All cells were washed with ice-cold phosphate-buffered saline (PBS) and then lysed for 1 hour on ice with RIPA buffer (Sigma-Aldrich, #R0278) containing 100 mM NaF, 4 mM Na-orthovanadate, 1 mM phenylmethylsulfonylfluoride (PMSF), 1 mM dithiothreitol (DTT), and 100 U/ml benzonase (Sigma-Aldrich). Primary bronchial ALI tissues were lysed for 2 hours on ice and with double concentration of benzonase. Samples were mixed with LDS Sample Buffer (Invitrogen), resolved by SDS-PAGE on 4-12 % Novex Bis-Tris precast gels (ThermoFisher Scientific) and analyzed by Western-blotting using a wet-transfer method (GenScript) and PVDF membranes (Life Technologies). The membranes were probed with mouse anti-human CFTR monoclonal antibodies n°596 or n°660 (Cystic Fibrosis Foundation, 1:2000) targeting NBD2 and NBD1 respectively, or mouse anti-human CFTR MAB3482 (Millipore, 1:200) targeting MSD1, and rabbit anti-GAPDH (#ab9485, Abcam, 1:5000) or rabbit anti-α/β-tubulin (Cell Signaling Technology, #2148; 1:2000) or mouse anti-ubiquitin (Santa Cruz, #P4D1, 1:500) or rabbit-anti-ubiquitin (Abcam, # #ab140601,1:200). Secondary antibodies were sheep horseradish peroxidase (HRP)–coupled anti-mouse IgG (Sigma-Aldrich, GENA931) and donkey HRP–coupled anti-rabbit IgG (Sigma-Aldrich, GENA934) were used at 1:5000 dilution. Membranes were treated with Western Lightning Enhanced Chemiluminescence Substrate (PerkinElmer), and the chemiluminescence signal was recorded and quantified using a chemiluminescence reader and the corresponding software (Fusion FX 7, Evolution Edge, Vilber).

### Immunofluorescence microscopy

CFBE-WT-CFTR and CFBE-F508del-CFTR cells were seeded at 10.000 cells/well onto FNC-coated 8-well chamber slides for a few hours and incubated with fixed concentrations of correctors or corresponding vehicle for 24 h. Then cells were washed once with PBS, fixed with 3 % paraformaldehyde (37°C, 10 min) in PBS, washed twice with PBS and then blocked with 10 % FBS, 0.1 % saponin in PBS (blocking buffer) for 1 h at room temperature. Cells were then stained with mouse monoclonal anti-CFTR antibody 570 (Cystic Fibrosis Foundation; 1:200 in above blocking buffer) and rabbit polyclonal anti-ZO-1 antibody (ThermoFisher Scientific #40-2200 1:200) for 2 h at room temperature, then washed three times in blocking buffer, followed by the addition of secondary antibodies in blocking buffer (goat-anti-rabbit-IgG-AlexaFluor488 [ThermoFisher Scientific #A11034] and goat-anti-mouse-IgG-AlexaFluor568 [ThermoFisher Scientific #A11031] both 1:200) and Hoechst 33342 (2 ug/mL from 10 mg/mL stock, ThermoFisher Scientific #H3570). After 30 min at RT, cells were washed 3x with blocking buffer, 2x with PBS and mounted. Images were taken with fixed settings using the Leica confocal microscope TCS SP5 and the 63x objective. Z-stacks of 10 slices (0.9 um distance) were taken and fused into one image by maximal intensity projection.

### Transepithelial short-circuit current measurements in reconstituted patient tissue

CFTR function was determined by the measurement of transepithelial short-circuit currents in reconstituted patient tissue using the using chamber. Tissues were either commercially available (MucilAir™, Epithelix) or prepared in-house from hAECs (Epithelix, Lonza). Measured tissues originated from six F508del homozygous patients (CF patient #1: CF-AB0609, female, 31 years-old; CF patient# 3: #28388-0000450918, Lonza, male, 25 years-old; CF patient#4: CF-MD0637 male, 22 years-old; CF patient #6: CF-MD0519, female, 33-years-old; CF patient #5: CF-MD0526, female, 16 years-old; CF patient #7: CF-MD0567, female, 39 years-old;), two F508del heterozygous patients (CF patient #8: CF-MD0638, F508del/2789+5G->A, male, 39 years-old; CF patient #9: CF-MD0487, F508del/2183AA->G, male, 24 years-old) and three non-CF donors (non-CF donor #5: WT-MD0810, female, 52 years-old; non-CF donor #2: WT-AB0834, male, 71 years-old; non-CF donor #4: CC-2540S-20TL256517, Lonza, female, 48 years-old). After co-incubation with correctors for 24 h in medium (MucilAir™ medium, Epithelix) supplemented with 60 % human serum (#4522, Sigma-Aldrich) at 37°C, 5% CO_2_, the inserts were mounted in Ussing chamber systems (EM-CSYS-8, Physiologic Instruments, San Diego CA, USA). The basal buffer was composed of (in mM) 110 NaCl, 1.2 CaCl_2_, 1.2 MgCl_2_, 2.4 Na_2_HPO_4_, 0.4 NaH_2_PO_4_, 25 NaHCO_3_, 5.2 KCl and NaOH to adjust the pH to 7.4, while the apical buffer contained (in mM) 120 Na-gluconate, 1.2 CaCl_2_, 1.2 MgCl_2_, 2.4 Na_2_HPO_4_, 0.4 NaH_2_PO_4_, 25 NaHCO_3_, 5.2 KCl and NaOH to adjust the pH to 7.4, the basal-to-apical chloride gradient facilitating the recording of chloride currents. During experiments, both buffers were maintained at 37 C and continuously gassed with a O_2_-CO_2_ gas mixture (95%-5%). Ag/AgCl electrodes filled with 3 M KCl were connected to both hemi-chambers to VCC MC8 voltage/current clamp amplifiers (Physiologic Instruments). Transepithelial short-circuit currents were recorded at 1 Hz sampling frequency using the Acquire and Analyze software version 2.3.8 (Physiologic Instruments) during a baseline period and upon sequential addition of 100 µM amiloride (Sigma-Aldrich), 5 µM forskolin (Sigma-Aldrich), 1 µM ivacaftor, 100 µM UTP (Sigma-Aldrich), and 20 µM Inh172 (Sigma-Aldrich). Offline analysis of short-circuit currents was performed with Excel and Prism 8 (GraphPad, San Diego, CA, USA), and then normalized against the surface of the inserts.

### Halide-sensitive-YFP quenching assay

U2OS-F508del-CFTR cells expressing Topaz-YFP F46L/H148Q/I152L were seeded at 20000 cells/well into 384-well black clear bottom plates in 40 µl/well growth medium (Mc Coy’s 5a, Gibco, 10% FBS, penicillin/streptomycin), containing the various CFTR correctors at the indicated concentrations. The cells were co-incubated with the compounds for 24 h (37°C, 5% CO_2_). The next day, plates were washed twice with 55 µl/well of PBS+ (PBS containing 0.9 mM Ca^2+^, and 0.5 mM Mg^2+^). PBS+ was fully removed and cells were supplemented with 15 µl PBS+. Then, 5 µl of 4x concentrated stocks of ivacaftor or vehicle in dilution buffer (PBS+, 0.1 µM forskolin, 0.2 % bovine serum albumin (fatty-acid free), pH 7.4) were added and incubated for 30 min in the dark. For experiments with I-172 (Sigma-Aldrich), the inhibitor was added to the mix with either ivacaftor or vehicle to reach a final concentration in cells of 20 µM. Then, plates were transferred to the FLIPR Tetra (fluorescence imaging plate reader, Molecular Devices: excitation 470-495 nm; emission: 526-585 nm), baseline fluorescence reading of the YFP signal was performed for 6 sec (10 x 0.6 second intervals) after which 25 µl of iodide buffer (137 mM NaI; 2.7 mM KCl; 1.5 mM KH_2_PO_4_; 8.1 mM Na_2_HPO_4_, 1 mM CaCl_2_; 0.5 mM MgCl_2_. pH 7.4) were added and fluorescence reading was continued for 70 seconds (50 x 0.6 second intervals; 20 x 2 second intervals) to assess YFP quenching through CFTR-mediated iodide influx. For analysis, fluorescence traces were aligned and normalized at the last time point before iodide addition (normalized fluorescence=1). Normalized fluorescence values obtained 20 seconds after the beginning of the assay were used to assess the degree of YFP quenching, i.e. CFTR function (see Figure S1J). For co-potentiation experiments, HEK-G551D-CFTR were seeded at 40000 cells/well into 384-well black clear bottom plates coated with 0.01% Poly-L-Lysine (Poly-L-Lysine, Sigma-Aldrich, #P8920, 0.1 % (w/v) diluted 1:10 in PBS-) in 40 µl /well of HEK growth medium. Cells were induced with 10 ng/mL tetracycline for 24 h (37°C, 5% CO_2_). The next day, plates were washed twice with 55 µl /well of PBS+ . PBS+ was fully removed and cells were supplemented with 15 µl PBS+. Ivacaftor in different concentrations or vehicle were mixed together with vehicle or correctors at the indicated concentrations (acute treatment). Then, 5 µl of 4x concentrated stocks of compound mix in dilution buffer (PBS+, 0.1 µM forskolin, 0.2 % bovine serum albumin (fatty-acid free), pH 7.4) were added and incubated for 10 min in the dark. Then, plates were read and analyzed as above.

### Metabolic pulse-chase and chase assays to determine CFTR folding efficiency and half-life

Stable isotope labeling using amino acids in cell culture (SILAC technology, ThermoFisher Scientific) was used. For pulse-chase assays to determine CFTR folding efficiency at the ER, CFBE-F508del-CFTR or CFBE-WT-CFTR were seeded at 100.000 cells/well into a FNC-coated 24-well plate in CFBE growth medium, and the next day incubated for 24 h with “SILAC light medium” (arginine- and lysine-free RPMI 1640 #A33823, 84 mg/L light L-arginine, # 89989, 146 mg/L light L-lysine, #88429 and 20 mg/L L-proline, # 89989 [to prevent a potential arginine-proline conversion]) supplemented with 10% dialyzed FBS. Cells were then washed once in PBS (37°C) and incubated for 1 hour with “starvation medium” (arginine-, lysine-free RPMI 1640, 1 % HSA) containing the indicated fixed concentrations of correctors. A 30-min metabolic pulse-labeling was then performed by replacing the medium with “SILAC heavy medium” [arginine- and lysine-free RPMI 1640, 20 mg/L L-proline, 84 mg/L heavy L-arginine (^13^C_6_^15^N_4_, #89990) and 146 mg/L heavy L-lysine (^13^C_6_^15^N_2_, #88209), 1% HSA] containing correctors. Cells were then washed with PBS and chased in “SILAC light medium” supplemented with 1 % HSA and correctors. This medium was kept for 1, 2.5, 4 and 6 h of chase time followed by cell lysis in RIPA buffer as described above. For chase assays to determine CFTR C-band half-life, CFBE-F508del-CFTR or CFBE-WT-CFTR were seeded as above and incubated for 24 h with “SILAC light medium” supplemented with 1% HSA and the indicated concentrations of correctors. Cells were then washed once in PBS (37^°^C) and incubated for 1 hour with “starvation medium”, followed by incubation with “SILAC heavy medium” both containing the correctors for 0.5, 1, 2.5, 4, 6, 8-9, 24 h of chase times followed by cell lysis. Cells for wash-out analysis, before addition of “SILAC heavy medium”, were washed twice in CFBE growth medium and once in PBS, and then incubated for the indicated chase times in “SILAC heavy medium” without correctors. For the chase assay in primary ALI tissues, fully differentiated tissues from CF-patient#2 and non-CF donors#2 and #3 were used and the experiment design was the same as for CFBE cells with all SILAC media supplemented with 1 % HSA and indicated correctors added directly at the basal side of ALI tissues. Samples were chased for 1, 2.5, 6, 8 and 25 h, followed by lysis in RIPA buffer as described above.

### Cross-linking with photo-activatable probes

U2OS-F508del-CFTR cells were seeded at 120.000 cells/well in a 24-well plate for a day and then incubated for 48 h with 1-2 µM of each probe (photo-activatable IDOR-5A, -6A, - 76B; non-photo-activatable controls IDOR-5, -6) diluted in the growth medium. Medium was then removed and the ice-cooled cell plate was UV irradiated (254 nm) at close distance (2 cm) without lid for 5 min (7 Watt, Camag). Cells were then lysed in RIPA buffer as described above. Control plates were not exposed and directly lysed.

### Mass spectrometry-based quantification of peptides

Cell lysates from pulse-chase, chase assays or cross-linking experiments were processed for LC/MS analysis. Samples were mixed with LDS sample buffer, resolved by SDS-PAGE on 3-8% tris-acetate precast gels (ThermoFisher Scientific), followed by protein staining with InstantBlue (Bio-Techne AG). CFTR protein bands were excised, cut in small pieces (1-2 mm^3^), moved to a Protein LoBind 96-well plate (Eppendorf) and destained with several alternating washes of freshly prepared solutions of 50 mM ammonium bicarbonate (Ambic), and 50 mM Ambic in 50 % acetonitrile (ACN), until the color was completely removed. Gel pieces were then dried under nitrogen and incubated for up to 1 h with a 5 mM TCEP-50 mM iodoacetamide in 50 mM Ambic solution for protein reduction and alkylation. After washes with 50 mM Ambic in 50 % ACN, gel pieces were dried again and incubated with a freshly prepared solution of 12.5 ng/µl trypsin (sequence grade, Promega) in 50 mM Ambic for 16-20 h at 37°C. Peptides were extracted from gels with one volume of 50 mM Ambic and two volumes of 5 % formic acid (FA) in 50 % ACN and collected in a new plate. Samples were concentrated to 1/3rd of their volume and resuspended in 0.1 % trifluoracetic acid (TFA) in H_2_O. Peptide solutions were loaded onto OASIS µElution 96-well plates (Waters), for desalting and enrichment. After several washes in 0.1% TFA, peptides were eluted under vacuum with 60 µl of 70 % ACN in 0.1% TFA, collected in a new LoBind plate, dried and resuspended in 30 µl of a 2 % ACN, 0.1% TFA solution.

The peptides were quantified on a Thermo Scientific Orbitrap Exactive HF-X instrument (ThermoFisher Scientific connected to an UltiMate 3000 RSLC nano system (ThermoFisher Scientific). The data was analyzed using Rmarkdown with R version 4.1.1 (2021-08-10) and plots were generated using the ggplot2 R package.

For metabolic pulse-chase and chase assays, typically 10 µl of the peptide solution was injected and quantified using a 30 min method. Peptides were loaded first on a Trap column (Acclaim PepMap 100 3 µm C18 column of 2 cm length and 75 µm inner diameter, ThermoFisher Scientific) and then injected on an analytical column (EasySpray 15 cm column 2 µm 100 Å C18 column with 75 µm inner diameter, Thermo Scientific) and separated using a 10 min gradient from 5 % Buffer B (80 % ACN and 0.1% FA in H_2_O) in Buffer A (0.1% FA in H_2_O) to 35% Buffer B. The eluting peptides were acquired using a MS1 spectra (200-1200 m/z, resolution of 60’000, automatic gain control target of 3e6, maximum injection time of 120 ms) followed by up to 39 MS2 spectra (isolation window 1.4 m/z with 0.2 m/z offset, 45’000 resolution, automatic gain control target of 2e5, maximum injection time of 86 ms) using a scheduled injection list for CFTR peptides, peptides of proteins for quality control assessment of the gel cutting and three iRT peptides (Biognosys, Schlieren).

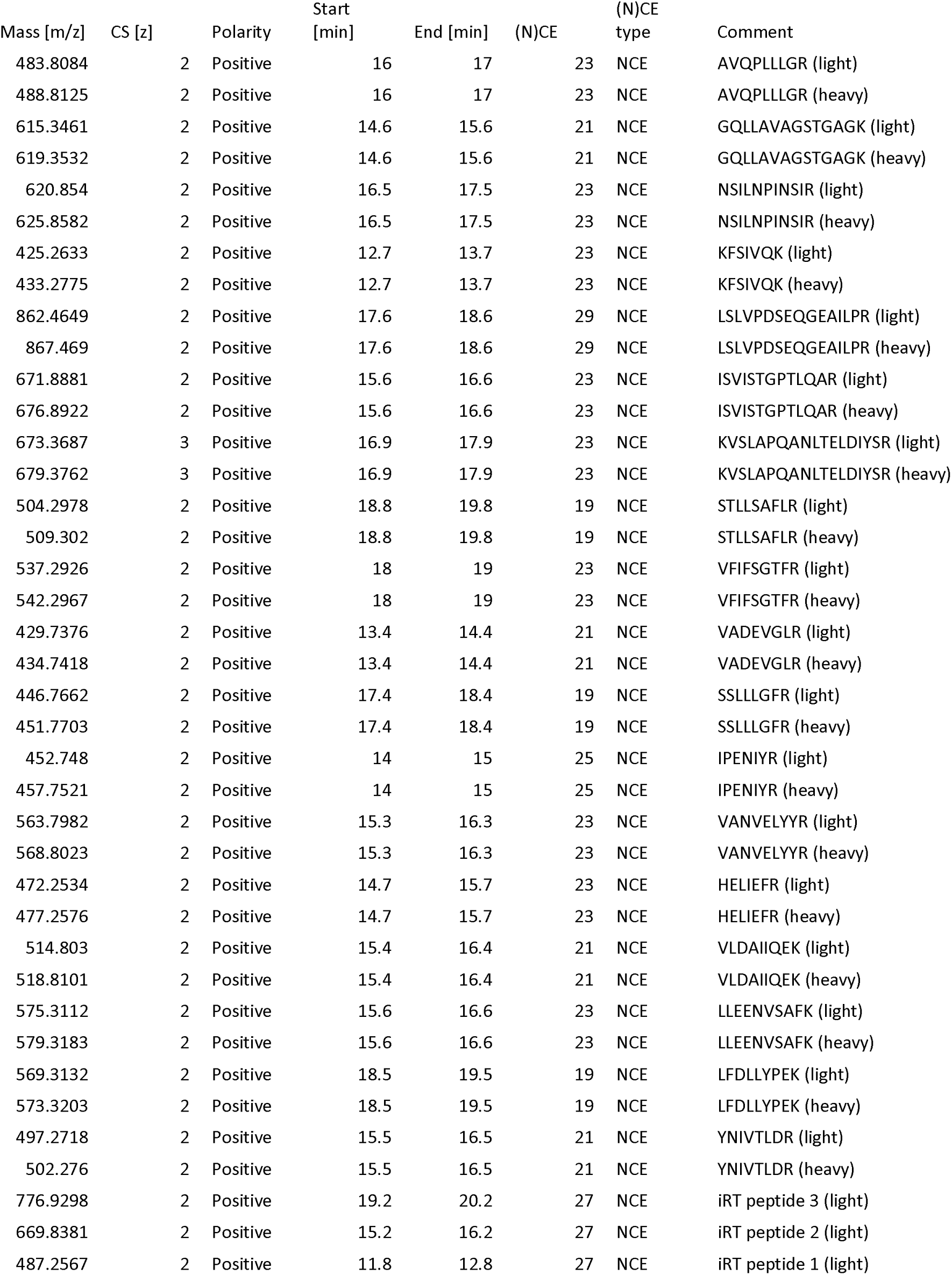

Data analysis of the signal acquired in parallel acquisition mode (PRM) was performed using Skyline (MacCoss Lab, University of Washington [40]). The signal for 4-6 fragments for each peptide were integrated by automatic peak-picking and manually controlled. The data was then exported to tab-delimited file for analysis using an in-house developed Rmarkdown script analyzing the total signal for all fragments for the 8 peptides (AVQPLLLGR, GQLLAVAGSTGAGK, NSILNPINSIR, LSLVPDSEQGEAILPR, ISVISTGPTLQAR, STLLSAFLR, VFIFSGTFR, VADEVGLR) from the CFTR protein. As a quality control (QC) step, we also analyzed QC peptides identified to co-migrate with the B- and C-band of CFTR (C-band: CLTC (theoretical molecular weight of protein sequence 192kDa) (VANVELYYR, HELIEFR) and SMC4 (147kDa) (VLDAIIQEK, LLEENVSAFK); B-band: MED23 (156 kDa) (LFDLLYPEK, YNIVTLDR) and LRPPRC (158 kDa) (SSLLLGFR, IPENIYR) to confirm precision of gel excision and separation of CFTR B- and C-bands (see Figure S3C).

Different peptides at the same concentration will give a different signal based on how easy they ionize, fragment etc., i.e. their response factor (see for an example Figure S3A). In this study, a relative response factor was calculated per experiment in the WT-CFTR cells for the light isotopologue relative to the AVQPLLGR peptide (see Figure S3B for an example). The signal of each peptide from CFTR was then normalized by their relative response factor. For the pulse-chase experiments, this normalized intensity was then further normalized to the maximal intensity observed in the B-band for the heavy isotopologue at time point 0.

For the cross-linking experiments, 10 µl of the gel-extracted peptide solution was injected and quantified using a 140 min method. Peptides were loaded first on a Trap column (Acclaim PepMap 100 3 µm C18 column of 2 cm length and 75 µm inner diameter, Thermo Fisher Scientific) and then injected on an analytical column (EasySpray 50 cm column 2 µm 100 Å C18 column with 75 µm inner diameter, Thermo Fisher Scientific) and separated using a 114 min gradient from 2 % Buffer B (80 % ACN and 0.1 % FA in H_2_O) in Buffer A (0.1% FA in H_2_O) to 35 % Buffer B. Masses from eluting peptides were acquired using a MS1 spectra (400-1200 m/z, resolution of 120’000, automatic gain control target of 3e6, maximum injection time of 20 ms) followed by up by 40 MS2 spectra in data-independent acquisition (DIA) mode from 400 m/z to 1000 m/z (isolation window 16 m/z with 100 m/z first mass, 30’000 resolution, automatic gain control target of 5e5, maximum injection time of 55 ms, normalized collision energy of 27). The data was then analyzed using a directDIA workflow in Spectronaut version 15 (Biognosys) using standard settings and a fasta file containing the mutated F508del-CFTR protein sequence. Transition-level data was exported and further filtered using in-house R scripts and mapDIA [41] and the intensities of 3-5 fragments per peptide are summed for analysis. The signal of each peptide was normalized to the total CFTR signal (summed signal of all CFTR peptides per sample) and to the average signal across all conditions of that respective peptide to account for the different response factors. Protein coverage in Figure S5E was displayed using the Protter [42].

### Chromatographic determination of probes concentrations upon UV light exposures

Stock solutions of IDOR-5A, IDOR-6A, IDOR-6B, IDOR-6 and IDOR-3 were prepared at 10 mM in DMSO. A series of twelve working standard solutions for calibration curve were prepared at concentrations from 250 nM to 1.22 nM by 2-fold serial dilution. 2 µM dilutions of the same compounds in 50 mM Tris buffer, exposed or not for 5 min to UV light as above, were diluted 10-fold with H_2_O/acetonitrile 50/50 v. Samples were then mixed and centrifuged at 3700 g for 15 min prior injection. The measurements were performed on a HPLC Nexera X2 HPLC (Shimadzu) coupled to an API 5500TM (AB Sciex) with positive ion electrospray ionization and operated in multiple reaction monitoring (MRM) mode. Data were collected and processed by Analyst® 1.6.2 and the chromatographic separation was carried out on a Acquity HSS T3 1.8 mm, 2.1x50 mm (Waters Corp) at 40 °C. The separation method consisted in a gradient elution of the mobile phase (0.1% formic acid in A: water and B: acetonitrile) as follows: 0 min 100% B, 98% B at 1.5 min and held for 0.6 min before re-equilibration. Total run time was 2.5 min, and all samples were analyzed with an injection of 1 mL. The source parameters were curtain gas (CUR), nitrogen: 20 psig, collision gas (CAD): 6 psig, ion source gas1: 20.0 psig; ion source Gas2: 20.0 psig; ion spray voltage (IS): 4500 V, turbo heater temperature (TEM): 500 °C; entrance potential (EP): 10 V. The electrospray ionization was operated in positive multiple reaction monitoring mode (MRM) after optimization according to standard procedure. Compound specific values of mass spectrometer parameters are listed in table below.

**Table:** Mass transitions, declustering potential and collision energy.

| Analyte | Q1<br>(m/z) | Q3<br>(m/z) | DP (V) | CE (V) |
| --- | --- | --- | --- | --- |
| IDOR-5A | 875 | 334 | 10 | 41 |
| IDOR-6A | 838 | 370 | 90 | 43 |
| IDOR-6B | 838 | 370 | 30 | 43 |
| IDOR-5 | 834 | 334 | 10 | 43 |
| IDOR-4 | 800 | 350 | 110 | 42 |

### CFTR mRNA quantification

CFBE-F508del-CFTR cells were seeded at 70.000 cells/well in a FNC-coated 48-well plate and incubated for 24 h with 2000 nM of indicated correctors or DMSO vehicle diluted in growth medium deprived of FBS, but supplemented with 1 % HSA. CFBE-WT-CFTR cells were treated with DMSO vehicle accordingly. Then cells were washed twice with PBS+ and lysed in RLT buffer (QIAGEN) for 10 min on ice. Lysates were fully homogenized with QIAshredder (#79656, QIAGEN) and total RNA was extracted with RNeasy Mini (#74106, QIAGEN) kit following manufacturer’s instruction. At the end a DNase treatment was performed with DNA-free kit (Invitrogen, AM1906) following manufacturer’s instruction. A reverse transcription (150 ng of total RNA per sample) with random hexamers was performed with the High capacity cDNA Reverse Transcription Kit (#4368814, ThermoFisher) for 2 hours at 37°C. 3µl of cDNA was mixed with 10 µl of TaqMan Fast Universal PCR Master Mix (2X) No AmpErase UNG (4367848, Life Tech.) containing the probe of interest (1:20) and a qPCR was run [20’’ at 95°C; 3’’ at 95°C – 30’’ at 60°C (40 cycles)] on a 7500 Applied Biosystems device. TaqMan (FAM-MGB) probes: *CFTR* (Hs00357011_m1) was used to detect the target gene; *18S* (Hs99999901_s1), *GUSB* (Hs00939627_m1) and *PPIA* (Hs04194521_s1) were used as reference genes for normalization. Reference gene-normalized CFTR mRNA expression was calculated using delta delta cT method with CFTR expression in vehicle-treated CFBE-F508del cells set to 1.

### 3D modeling

All computational calculations were performed with the phosphorylated structure of human CFTR (PDB code 6MSM) and F508del-CFTR (PDB code: 8EIQ). The two Mg-ATP complexes were kept, and the membrane was positioned using the OPM database [43]. F508 was deleted and the immediate surrounding was subject to minimization using Maestro (https://www.schrodinger.com/products/maestro). Docking and Molecular Dynamic Simulations were conducted with Glide XP [44] and Desmond [45] (Schroedinger suite), respectively. MD simulations used F508del-CFTR embedded in a POPC lipid bilayer. The overall system was solvated (TIP3P) and neutralized with 150 nM NaCl. Each simulation (250 ns) was conducted at 300K using the default parameters.

### Statistics

For evaluation of corrector efficacies on second-site suppressor mutants in HEK cells, a one-way ANOVA test with Turkey’s multiple comparison post-test was done between the three different mutants within each compound treatment. The same test was performed to evaluate corrector efficacies in binding sites double or triple mutants, between the four different corrector treatments within every mutant. Statistical tests were done with GraphPad Prism 8.

To identify peptides that showed lower signal upon cross-linking, the normalized value for all >80 quantified peptides of CFTR was statistically compared across conditions using pair-wise testing for a loss of signal (i.e. one-sided t-test), using R software 4.1.1. We compared the UV-excited conditions of samples treated with each compound to their respective non-exposed controls. The signal of each cross-linkable compound after UV-excitation was also compared to the respective UV-excited non-crosslinkable parent compound (IDOR-5A vs IDOR-5 and IDOR-6A or IDOR-6B vs IDOR-6). To control for multiple testing of the about 80 peptides in each comparison, a multiple-testing correction using Benjamini Hochberg was performed (p.adj). With these stringent statistical criteria only two peptides LFFSWTR and LFFSWTRPILR showed a statistically significant difference in signal after multiple testing correction (p.adj = 2.0·10^-4^ and p.adj = 1.2·10^-5^ and respectively) in the comparison of IDOR-6A vs IDOR-6. No other peptide in no other tested comparison showed an adjusted p-value below 0.05 (Figure S5F). The results for these two peptides without multiple testing correction showed that there was also a statistical different signal between IDOR-6A treated samples after and without UV excitation (p = 7.5·10^-4^ and p = 1.4·10^-3^ respectively) but not in any other comparison.

## ACKNOWLEDGEMENTS

We thank Dr. J.P. Clancy (University of Alabama, Birmingham) for providing CFBE41o-cells stably expressing human WT-CFTR or F508del-CFTR mutant, Dr. J. Riordan and Dr. Martina Gentzsch (University of North Carolina at Chapel Hill) as well as the Cystic Fibrosis Foundation for providing CFTR antibodies 596, 660 and 570. At Idorsia Pharmaceuticals, we thank Dr. Alexia Chavanton-Arpel, for the development of Ussing chamber experiments. We also thank, Diego Freti for mRNA quantification and François Le Goff for HPLC analysis. For expert synthesis support, we gratefully thank Christian Barten, Aude Bauer, Eser Ilhan, Thierry Kimmerlin, Pauline Ligibel, Cristian Linder, and Sylvain Regeon.

## Funding

No external funding was received.

## Author contributions

Conceptualization: JG, VM

Methodology: VM, LM, PB, CK, JG, RS, JK, CF, JTW, CB

Investigation: VM, LM, PB, CK, AB, JK, CF, JG Visualization: JG, PB, FC, AB, VM Writing—original draft: VM, JG Writing—review & editing: ON, EE, PB, JG

## Competing interests

All authors are employees of Idorsia Pharmaceuticals, Ltd. JG, JTW, CB are authors of patent WO2022194399A1 describing some of the type-IV correctors used in this study.

## Data and materials availability

All data are available in the main text or the supplementary materials.

---

**Fig. S1.**
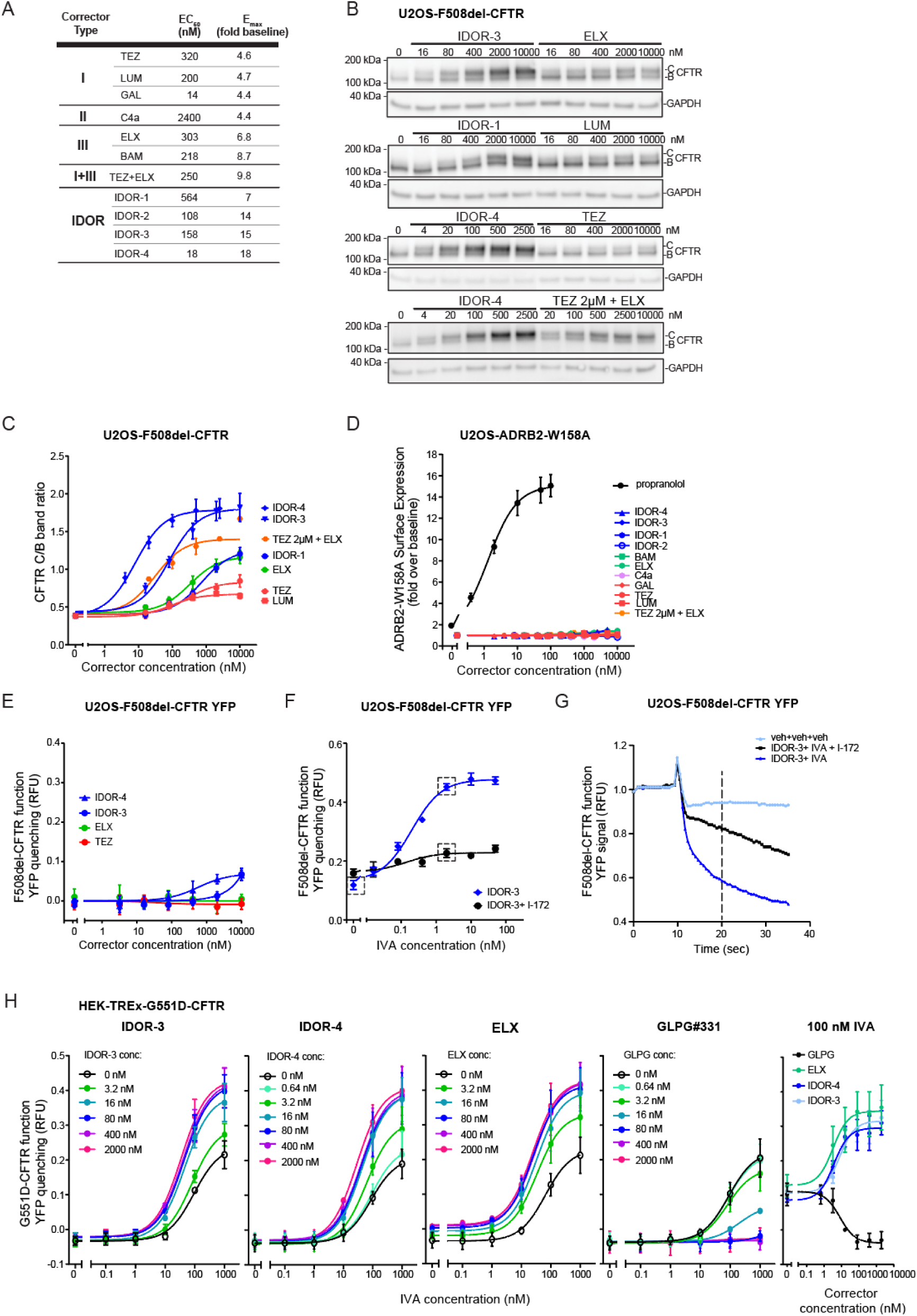
Related to Fig. 1: Type-IV correctors restore F508del-CFTR trafficking to the cell surface and behave additively with type-I, type-II, and type-III correctors. **(A)** EC_50_ and E_max_ values deduced from the corrector concentration response curves shown in Figure 1A. **(B)** Immunoblot analysis of CFTR and GAPDH expression in U2OS-F508del-CFTR cells treated over-night with different concentrations of the indicated correctors. Representative images of n=7 independent experiments. **(C)** Quantification of the CFTR C-band intensity normalized for the B-band (n=2 for TEZ+ELX combination, n=7 for all other compounds). **(D)** ADBR2^W158A^ surface expression in U2OS cells upon over-night treatment with dilution series of the indicated correctors or adrenergic receptor ligand propranolol (n=3 for TEZ+ELX combination, n=5 for all other compounds). **(E-F)** F508del-CFTR function (Topaz-YFP^F46L.H148Q.I152L^quenching assay) after 24 h treatment of U2OS-F508del-CFTR-YFP cells with: E) Concentration response of IDOR-3, IDOR-4, TEZ or ELX in the acute presence of 0.1 µM forskolin (n=2); F) 2000 nM of IDOR-3 in presence or absence of different concentrations of IVA and with or without 20 µM of CFTR-inhibitor-172 (n=3). Enclosed in dashed squares are the conditions shown in (G). **(G)** Examples of raw YFP quenching traces of one of the experiments shown in I (veh or 2000 nM IDOR-3 + 2 nM IVA or 2000 nM IDOR-3 + 2 nM IVA + 20 µM I-172). Fluorescence values obtained 20 seconds from the beginning of the measurement were used to assess the degree of YFP quenching. **(H)** G551D-CFTR function (YFP quenching assay) in HEK-TREx-G551D-CFTR-YFP cells exposed to acute treatment (10 min) with different concentrations of IVA combined with different concentrations of IDOR-3, IDOR-4, ELX or GLPG #331 (n=2). The right panel replots the data as corrector concentration-response curves for the 100 nM IVA condition. Data in (C), (D), (E), (F) and (H) are means ± SEM of the indicated number of independent experiments.

**Fig. S2.**
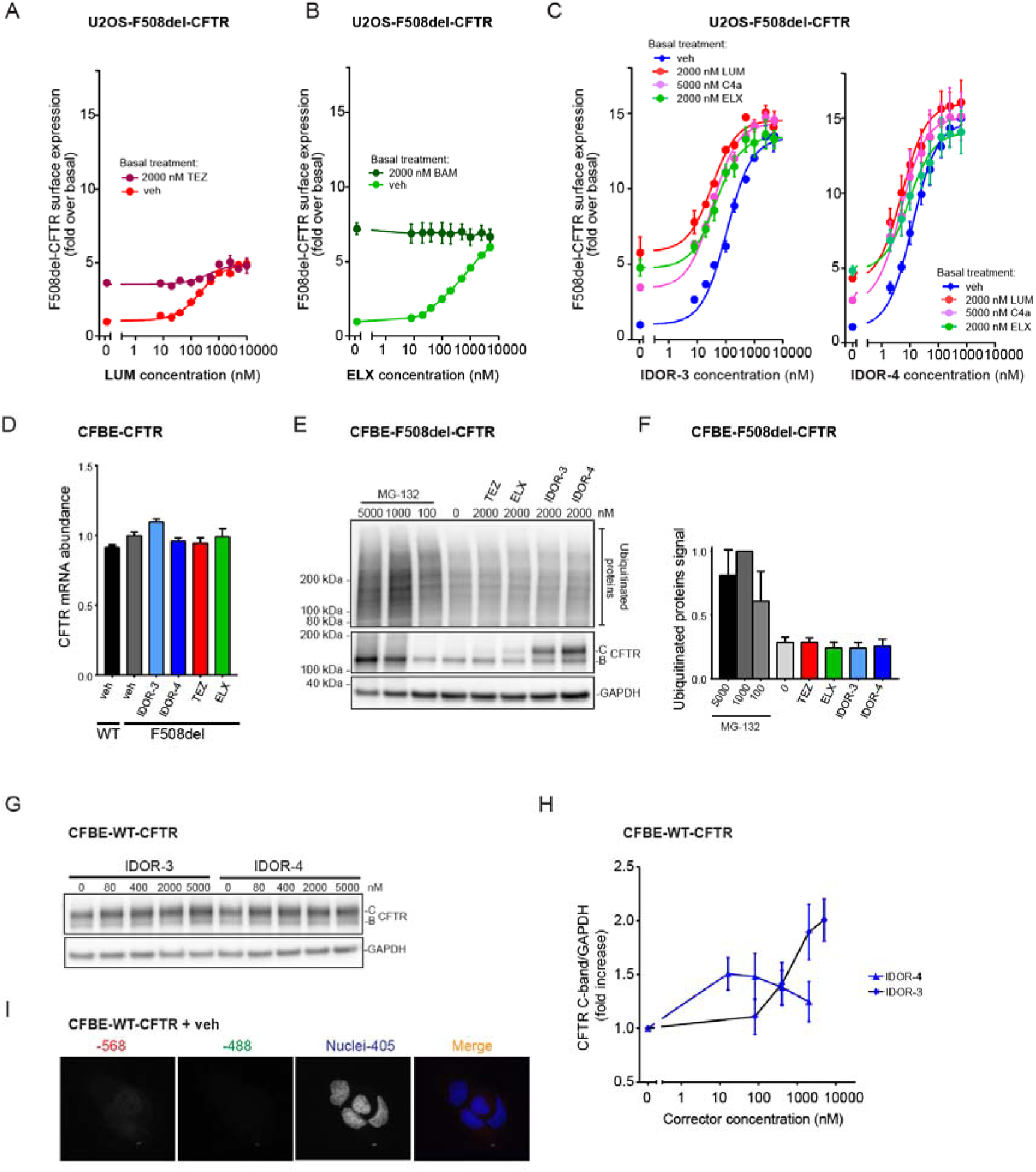
Related to Fig. 1: Type-IV correctors restore F508del-CFTR trafficking to the cell surface and behave additively with type-I, type-II, and type-III correctors. **(A-C)** CFTR surface expression in U2OS-F508del-CFTR cells after over-night treatment with: A) different concentrations of LUM in presence of vehicle or a maximally effective concentration of TEZ (n=3). B) different concentrations of ELX in presence of vehicle or a maximally effective concentration of BAM (n=4). C) different concentrations of IDOR-3 or IDOR-4 in presence of vehicle or a maximally effective concentration of the indicated other correctors (n=3); **(D)** Quantification of CFTR mRNA expression in CFBE41o-cells expressing WT-CFTR (n=2) or F508del-CFTR after 24 h of treatment with DMSO (n=3) or 2000 nM of the indicated correctors (n=2). **(E)** Immunoblot analysis of levels of ubiquitinated proteins, CFTR and GAPDH in CFBE-F508del-CFTR cells after 24 h of treatment with DMSO, MG-132 or the indicated correctors. Representative images (n=3). **(F)** Quantification of ubiquitinated proteins in (E), normalized for GAPDH (n=3). **(G)** Immunoblot analysis of WT-CFTR-expressing CFBE41o-cells treated for 24 h with different concentrations of the indicated correctors and probed for CFTR and GAPDH. Representative images (n=3). **(H)** Quantification of the CFTR C-band intensities in (G) normalized for GAPDH (n=3). **(I)** Representative control immunofluorescence pictures for Figure 1G showing CFBE-WT-CFTR cells treated with DMSO for 24 h and stained for cell nuclei in blue and only secondary antibodies (red and green channels). Data in (A), (B), (C), (D), (F) and (H) are mean ± SEM of the indicated number of independent experiments.

**Fig. S3.**
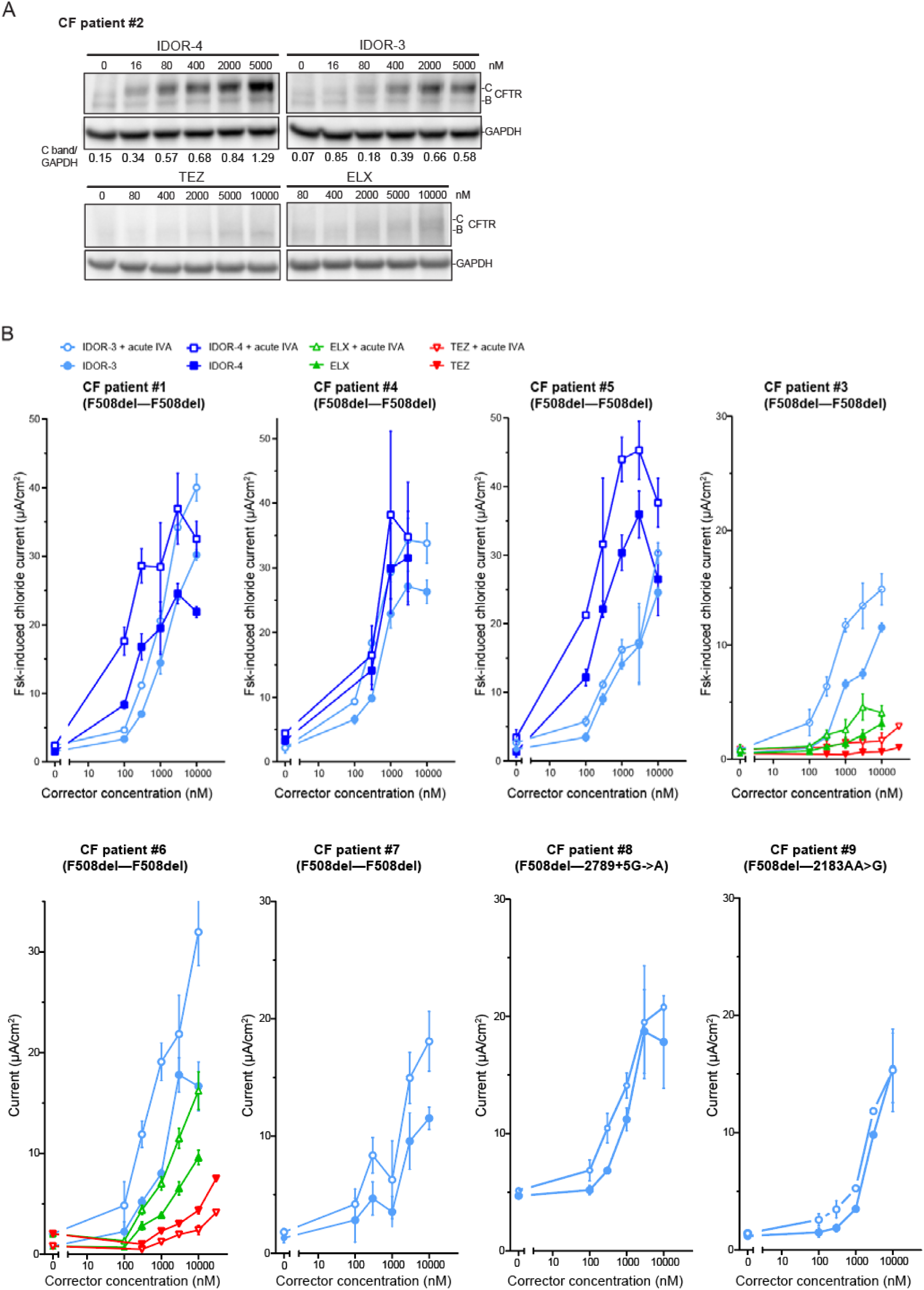
Related to Fig. 2: Type-IV correctors rescue F508del-CFTR trafficking and function in reconstituted cystic fibrosis bronchial epithelium. **(A)** CFTR expression (immunoblotting) in reconstituted human bronchial epithelium of CF patient #2 (F508del-CFTR homozygous) after a 24-h treatment with the indicated correctors. Representative images (n=2). The CFTR C-band/GAPDH intensity ratio is represented below every lane (for TEZ and ELX correction the ratio could not be calculated because the signal was too weak). **(B)** Measurement of transepithelial short-circuit currents in Ussing chamber in reconstituted epithelium of 8 CF patients (6 homozygous and two heterozygous for F508del-CFTR), treated over-night with vehicle or IDOR-3, IDOR-4, TEZ or ELX dilution series, in presence or absence of acute 1 µM IVA (at least 2 independent measurements/patient). Data in (B) are mean ± SEM of the indicated number of independent experiments.

**Fig. S4.**
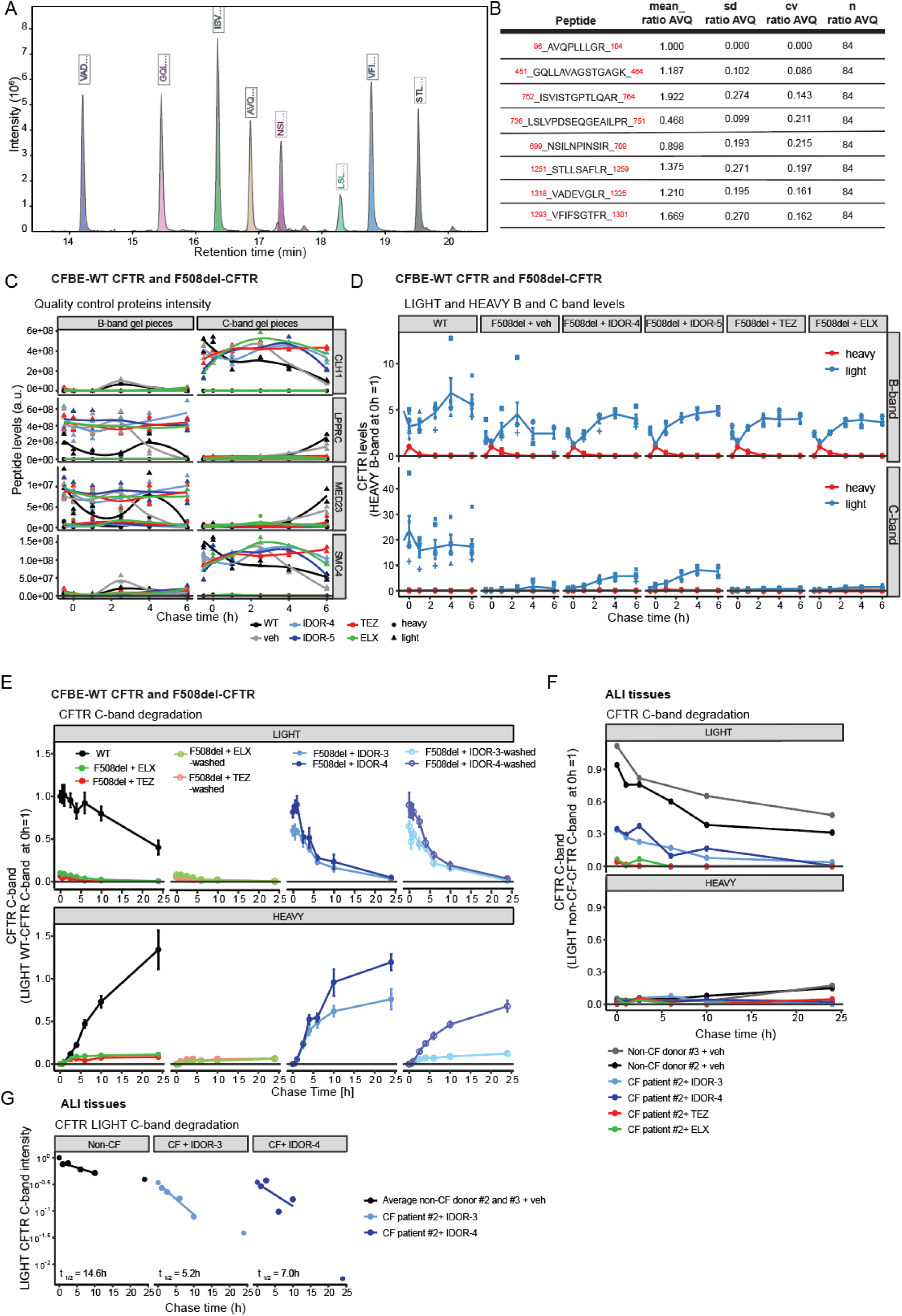
Related to Fig. 3: Type-IV correctors restore F508del-CFTR folding efficiency in the endoplasmic reticulum beyond wildtype levels. **(A)** Representative chromatogram showing the elution of the 8 peptides quantified for CFTR by mass spectrometry. **(B)** Representative normalization ratio of each peptide. All peptides are normalized to each other to account for the different response factor using the mean_ratio_AVQ which is the mean ratio of intensity between the respective peptide and the peptide AVQPLLLGR. **(C)** Average peptide intensities of heavy-labeled (circles) and light-labelled (triangles) peptides of proteins predicted to co-migrate with CFTR-B band (LPPRC and MED23) or C-band (CLH1 and SMC4). Samples originate from pulse-chase experiments with CFBE cells expressing WT-CFTR or F508del-CFTR, pulse-labeled for 30 min with heavy amino acids and chased with light amino acids for the indicated time periods. During the whole assay F508del-CFTR cells were treated with DMSO (n=5) or 2 µM IDOR-3 (n=4), 2 µM IDOR-4 (n=2), 10 µM TEX or 10 µM ELX (n=3), indicated with different colors. **(D)** Average peptide intensities of heavy-labeled (red lines) and light-labeled (blue lines) CFTR B-(upper graphs) and C-(lower graphs) bands in CFBE cells treated as described under C). Peptide intensity per time point in each treatment is normalized to maximal intensity of heavy-labeled-B band at 0h chase time. **(E)** Average peptide intensities of light and heavy CFTR C-band in cells incubated for 24 h with light amino acids in the presence of DMSO (n=4) or 2 µM IDOR-3 or 1 µM IDOR-4 or 10 µM TEZ or 10 µM ELX (n=2 each) and then chased with heavy amino acids. Correctors were either kept during the chase period or thoroughly washed-out before the chase. Per experiment, peptide intensities at the different time points were normalized to the intensity of light C-band of CFTR WT at the beginning of the chase. **(F)** Peptide intensities of light- and heavy-labeled CFTR C-band in ALI tissues from non-CF donors #2 and #3 and CF patient #2, incubated for 24 h with light amino acids and DMSO or 2 µM IDOR-3, 1 µM IDOR-4, 10 µM TEZ or 10 µM ELX and then chased for the indicated time points with heavy amino acids. Per experiment, peptide intensities were normalized to the maximal intensity of the light-labeled C-band of WT-CFTR at the beginning of the chase (n=1). **(G)** Calculation of the light-labeled CFTR C-band half-life in ALI tissues from non-CF donors #2 and #3 and CF patient #2 corrected with IDOR-3 and IDOR-4, determined by linear regression of the logarithmic data using time points up to 10 h (n=1). Data in (C), (D) and (E) are mean ± SEM of the indicated number of experiments.

**Fig. S5.**
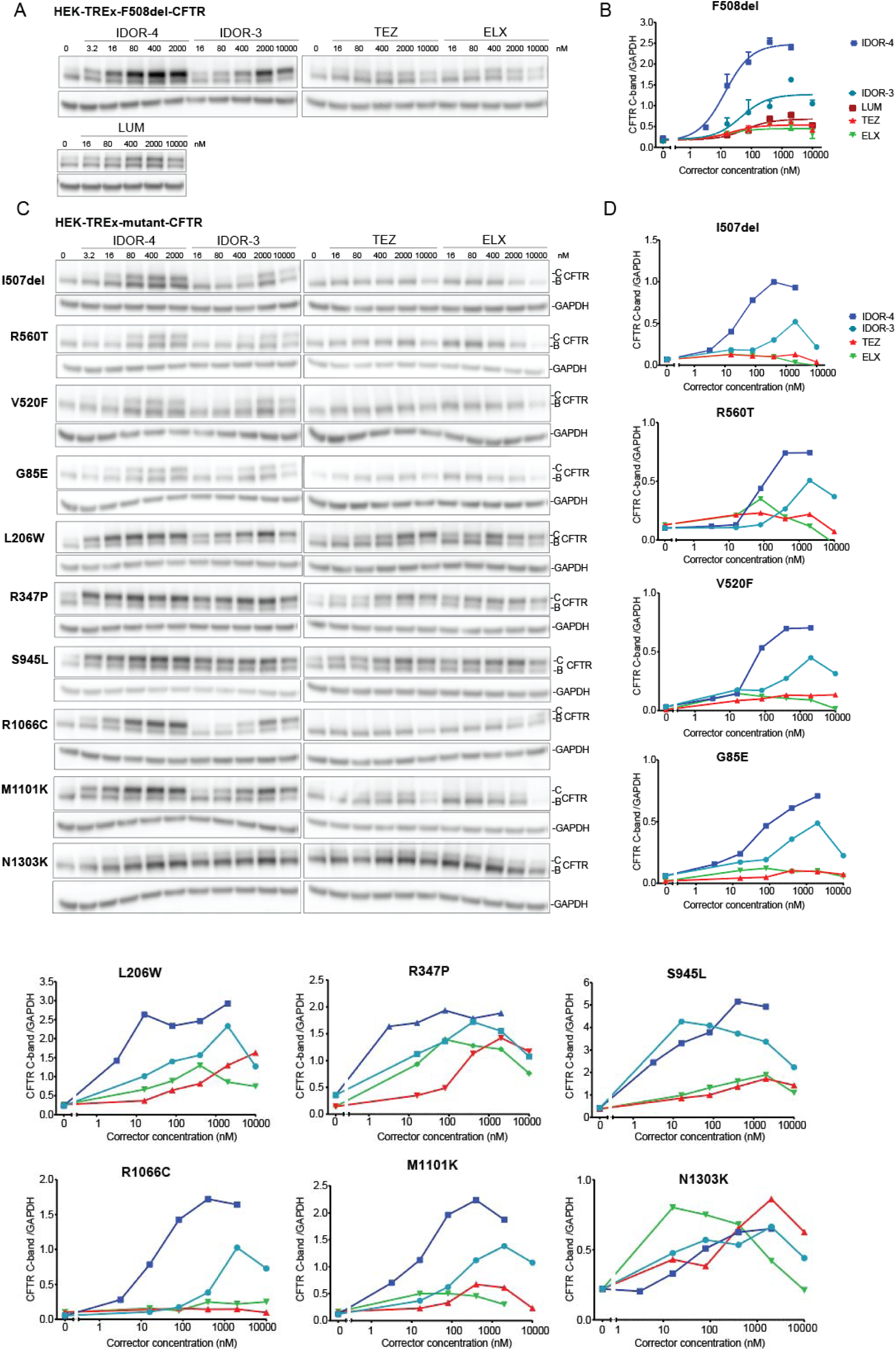
Related to Fig. 4 and 6: Type-IV correctors address non-F508del CFTR folding mutations. **(A and C)** Representative immuno-blot images of CFTR and GAPDH in HEK-TREx cells expressing 11 different CF-causing CFTR folding mutations and treated for 48 h with the indicated corrector concentrations (n=2 for A, n=1 for C). (**B and D)** CFTR C-band intensities normalized for GAPDH (n=2 for A, n=1 for C). Data in (B) are means ± SEM of the indicated number of independent experiments. In (D) values are from one experiment.

**Fig. S6.**
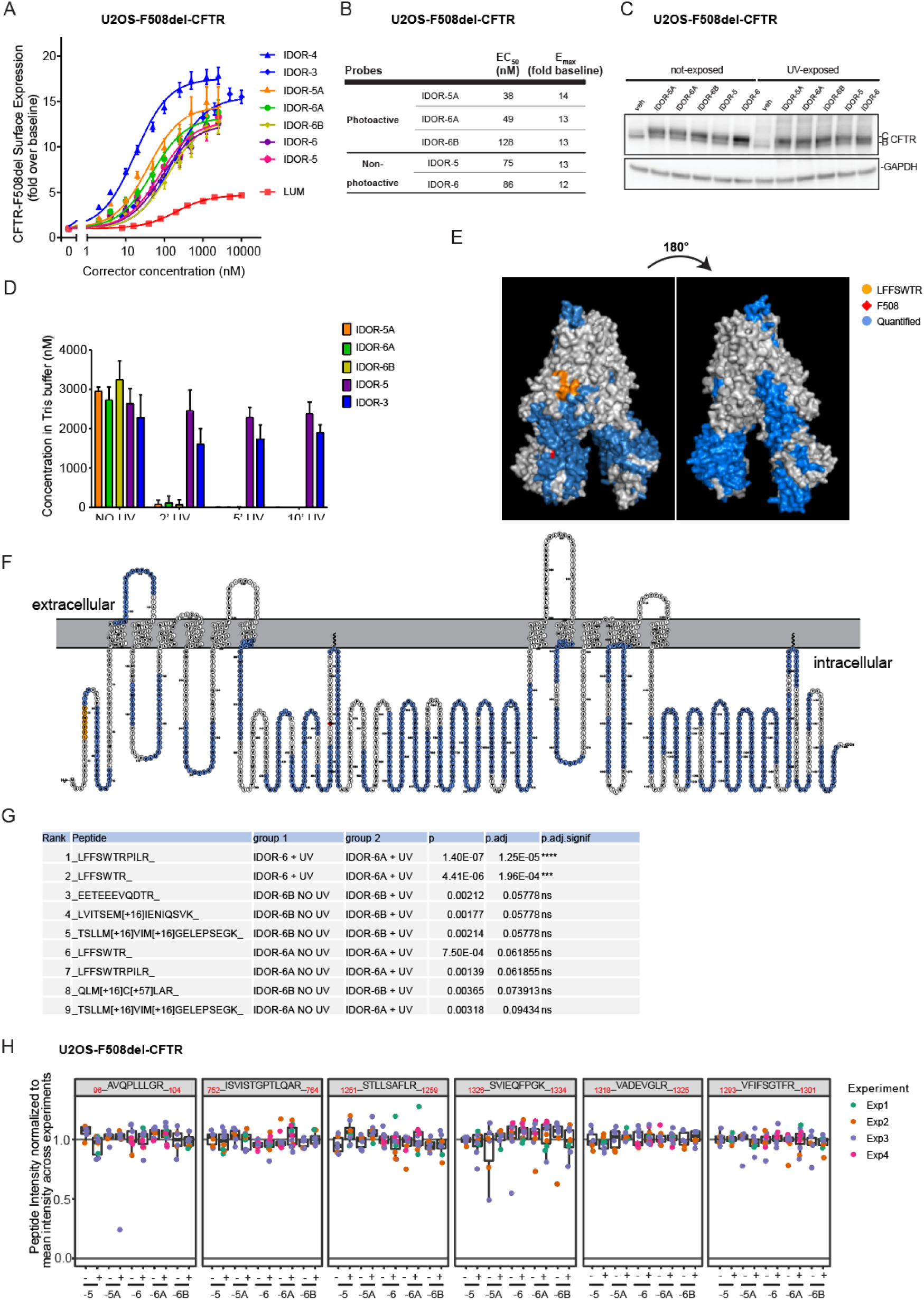
Related to Fig. 5: Type-IV correctors bind to the CFTR lasso domain. **(A)** F508del-CFTR cell surface expression in U2OS cells after overnight treatment with different concentrations of the indicated correctors (n=3 for each probe). **(B)** EC_50_ and E_max_ values of the three photoactivatable probes and the two parent macrocycles determined in (A). **(C)** CFTR and GAPDH expression levels (immuno-blotting) in U2OS F508del-CFTR cells treated for 48 h with 2 µM of the indicated probes and exposed or not to UV light for 5 min. Representative images (n=2). **(D)** Concentration change of the indicated probes diluted in 50 mM Tris buffer with increasing UV exposure times (n=6). **(E and F)** 3D and 2D view of the CFTR protein showing in blue all 73 peptides detected in the crosslinking approach in at least two experiments, corresponding to 53 % of the whole protein. Peptide15-21 is shown in yellow and F508 in red. **(G)** Table showing the first 9 of all identified peptides, ranked from the lowest adjusted p-value after one-sided t-test and multiple-testing correction using Benjamini Hochberg test (see Methods). _15_LFFSWTR_21_ and _15_LFFSWTRPILR_25_ were the only peptides statistically significantly reduced in any comparison and were different for samples treated with IDOR-6A versus the control probe IDOR-6 (after UV exposure). The non-adjusted p-value for these two peptides (without multiple testing correction) showed also a statistically significant difference between IDOR-6A treated samples after and without UV excitation. **(H)** Abundance of six representative CFTR peptides showing no specific effect after UV exposure or treatment with the six different probes (n=3/4). Data in (A) are mean ± SEM of the indicated number of experiments. Data in (D) are mean ± SD of 6 experimental replica. Scatter plot in (H) results from at least 3independent experiments with colored dots representing replica values. Per experiment, the intensity level of each peptide was normalized to the total CFTR signal in that sample (summed signal of all CFTR peptides).

**Fig. S7.**
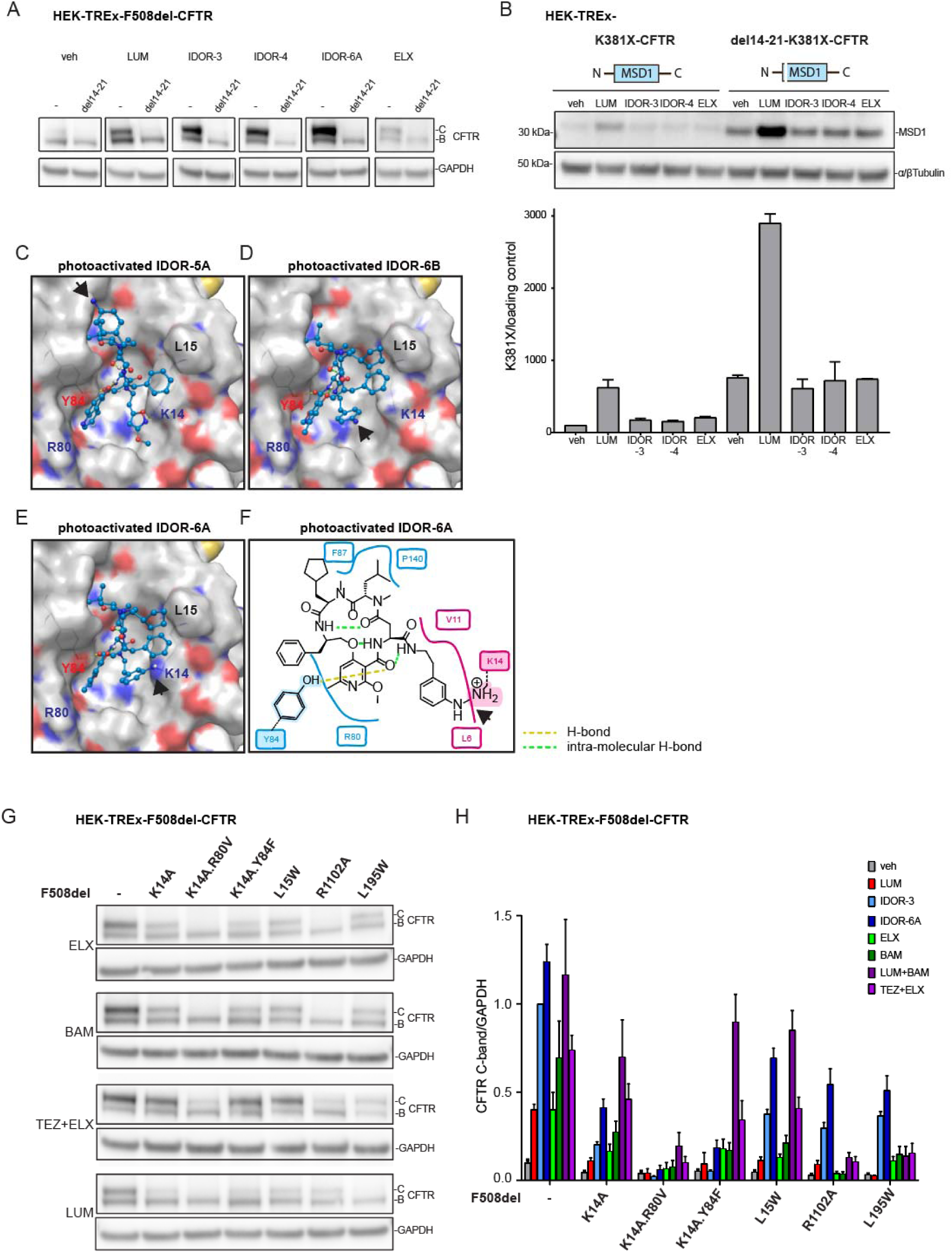
Related to Fig. 5: Type-IV correctors bind to the CFTR lasso domain. **(A)** CFTR and GAPDH expression by immuno-blotting in HEK-TREx cells expressing F508del-CFTR with or without additional deletion of the amino acids 14-21 and treated for 40 h with 2 µM LUM or 2 µM IDOR-3 or 0.4 µM IDOR-4 or 2 µM IDOR-6A or 0.4 µM ELX (n=2). **(B)** Levels of truncated CFTR construct K381X with or without amino acid 14-21 deletion expressed in HEK-TREx cells after 40 h of treatment with vehicle or 2 µM LUM or 2 µM IDOR-3 or 0.4 µM IDOR-4 or 0.4 µM ELX. Detection antibody targets MSD1. Intensities of CFTR fragment bands are shown in the graph below, normalized for the loading controls (n=2). **(C-E)** 3-dimensional views of photo-activated IDOR-5A (C), IDOR-6B (D) and IDOR-6A (E) bound to the cavity as determined by docking studies. Polar amino acid side chains are indicated in blue, red and yellow. H-bonds between the probes and CFTR are indicated as yellow dashed lines. Intramolecular H-bonds are depicted as green dashed lines. Black arrowheads point to the aryl-nitrene (activated intermediate) in each probe. **(F)** 2-dimensional view of IDOR-6A with likely cross-linking of the aryl-meta-nitrene to the K14 amine. **(G)** Site-directed mutagenesis of key amino acids in the proposed type-IV corrector binding site and effect on corrector activities: F508del-CFTR constructs containing one or two additional mutations in the site (K14A, K14A.R80V, K14A.Y84F or L15W) or the ELX binding site (R1102A) or the LUM binding site (L195W) were expressed in HEK-TREx cells then treated for 40 h with 2 µM of each indicated corrector. Representative images (at least 4 independent experiment per mutant and per corrector treatment). **(H)** Quantification of F508del-CFTR C-band intensities in (F) normalized for GAPDH (n=4-10). Data in (B) and (H) are mean ± SEM of the indicated number of experiments.

