## Supplementary Figures for "A Novel, Uniquely Efficacious Type of CFTR Corrector with Complementary Mode of Action"

**SUPPLEMENTAL FIGURES**

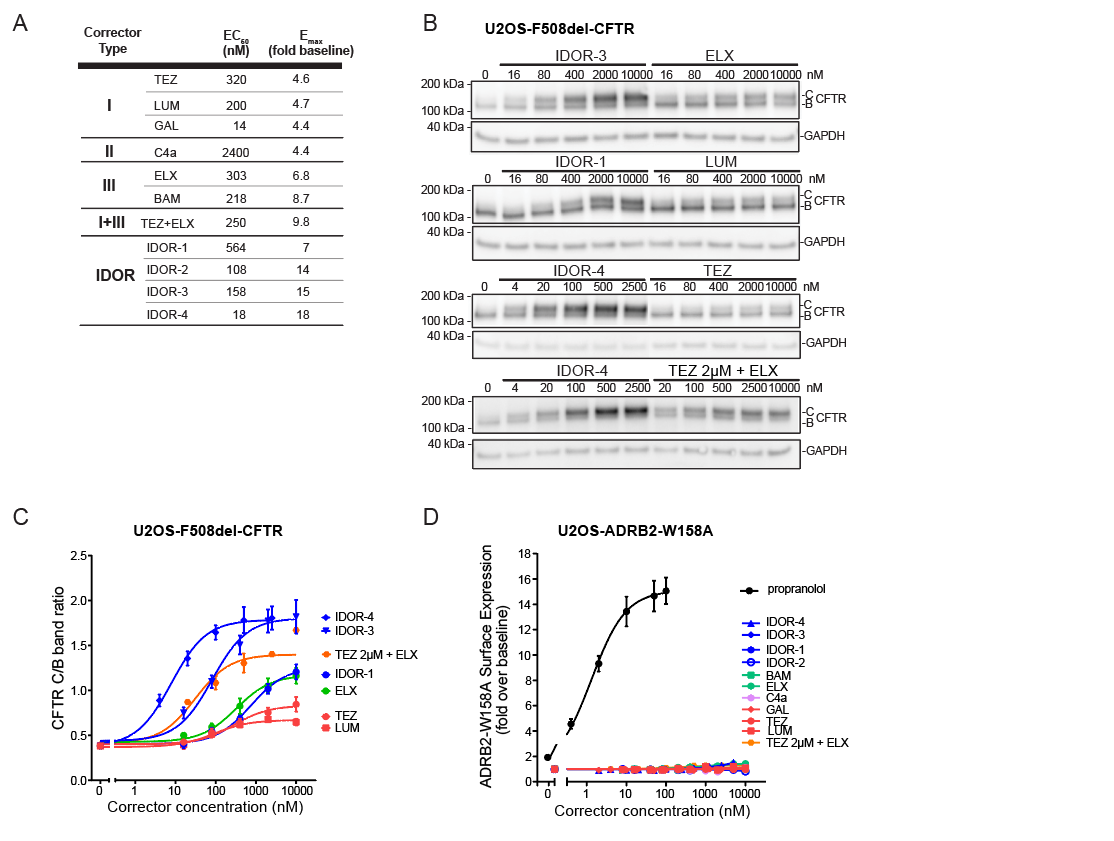

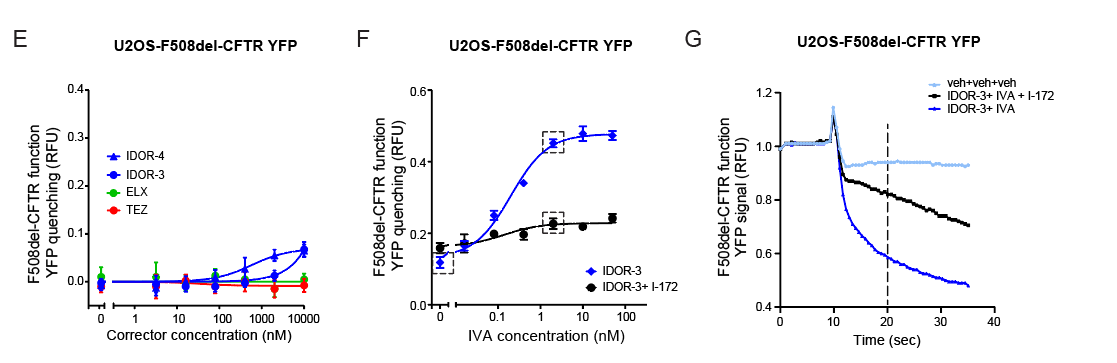

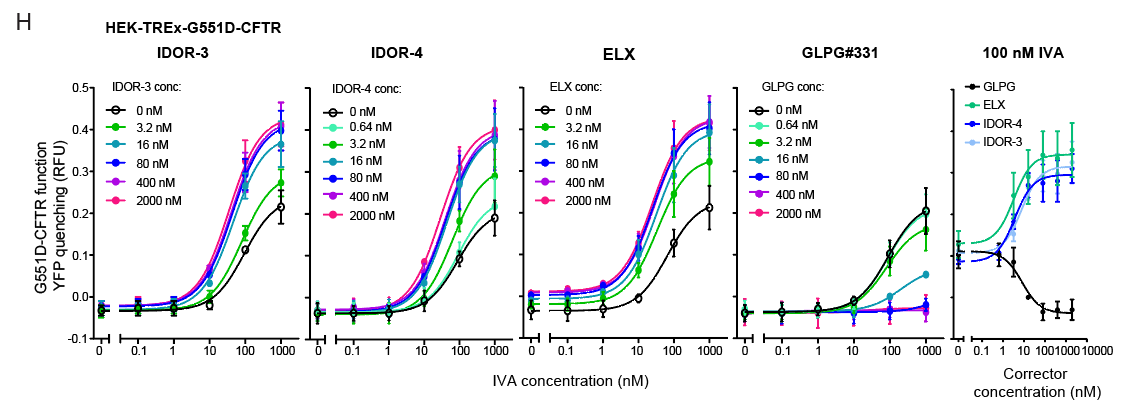
***Fig. S1. Related to Fig. 1: Type-IV correctors restore F508del-CFTR trafficking to the cell surface and behave additively with type-I, type-II, and type-III correctors. (A)*** *EC_50_ and E_max_ values deduced from the corrector concentration response curves shown in Figure 1A.* ***(B)*** *Immunoblot analysis of CFTR and GAPDH expression in U2OS-F508del-CFTR cells treated over-night with different concentrations of the indicated correctors. Representative images of n=7 independent experiments.* ***(C)*** *Quantification of the CFTR C-band intensity normalized for the B-band (n=2 for TEZ+ELX combination, n=7 for all other compounds).* ***(D)*** *ADBR2^W158A^ surface expression in U2OS cells upon over-night treatment with dilution series of the indicated correctors or adrenergic receptor ligand propranolol (n=3 for TEZ+ELX combination, n=5 for all other compounds).* ***(E-F)*** *F508del-CFTR function (Topaz-YFP^F46L.H148Q.I152L^quenching assay) after 24 h treatment of U2OS-F508del-CFTR-YFP cells with: E) Concentration response of IDOR-3, IDOR-4, TEZ or ELX in the acute presence of 0.1 µM forskolin (n=2); F) 2000 nM of IDOR-3 in presence or absence of different concentrations of IVA and with or without 20 µM of CFTR-inhibitor-172 (n=3). Enclosed in dashed squares are the conditions shown in (G).* ***(G)*** *Examples of raw YFP quenching traces of one of the experiments shown in I (veh or 2000 nM IDOR-3 + 2 nM IVA or 2000 nM IDOR-3 + 2 nM IVA + 20 µM I-172). Fluorescence values obtained 20 seconds from the beginning of the measurement were used to assess the degree of YFP quenching.* ***(H)*** *G551D-CFTR function (YFP quenching assay) in HEK-TREx-G551D-CFTR-YFP cells exposed to acute treatment (10 min) with different concentrations of IVA combined with different concentrations of IDOR-3, IDOR-4, ELX or GLPG #331 (n=2). The right panel replots the data as corrector concentration-response curves for the 100 nM IVA condition. Data in (C), (D), (E), (F) and (H) are means ± SEM of the indicated number of independent experiments.*

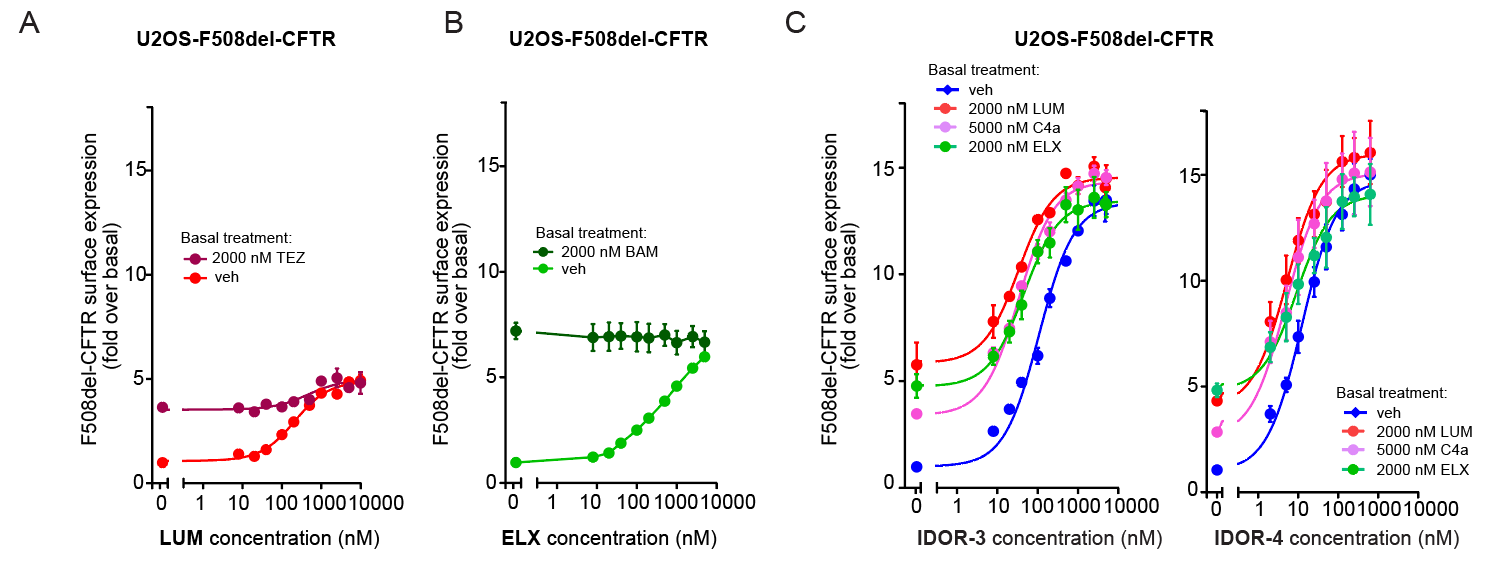

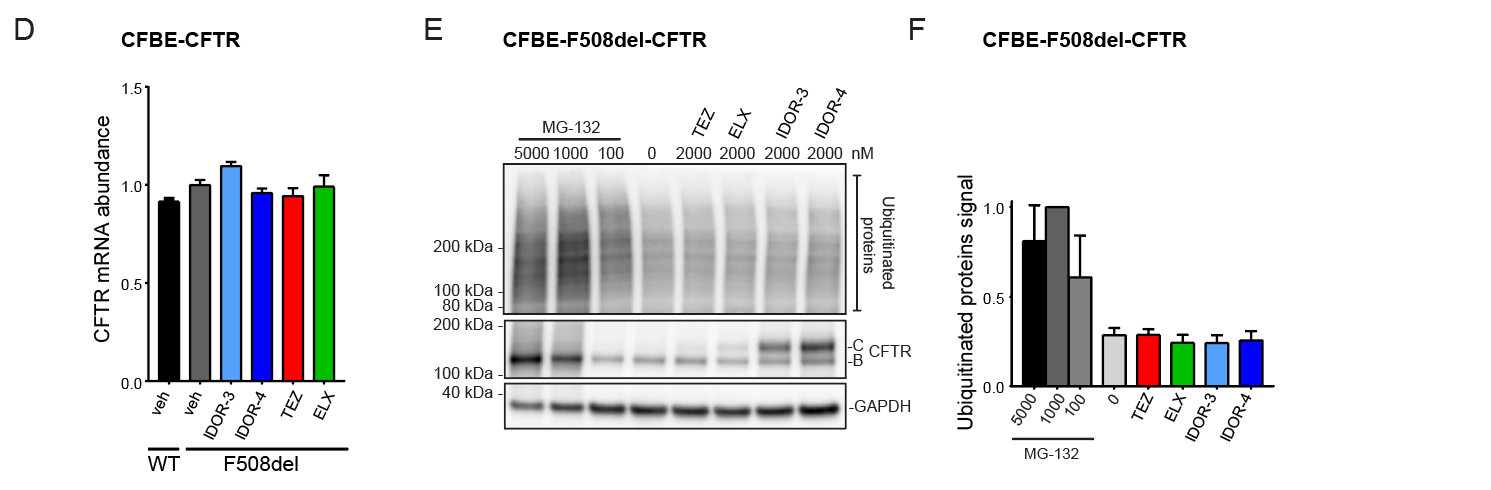

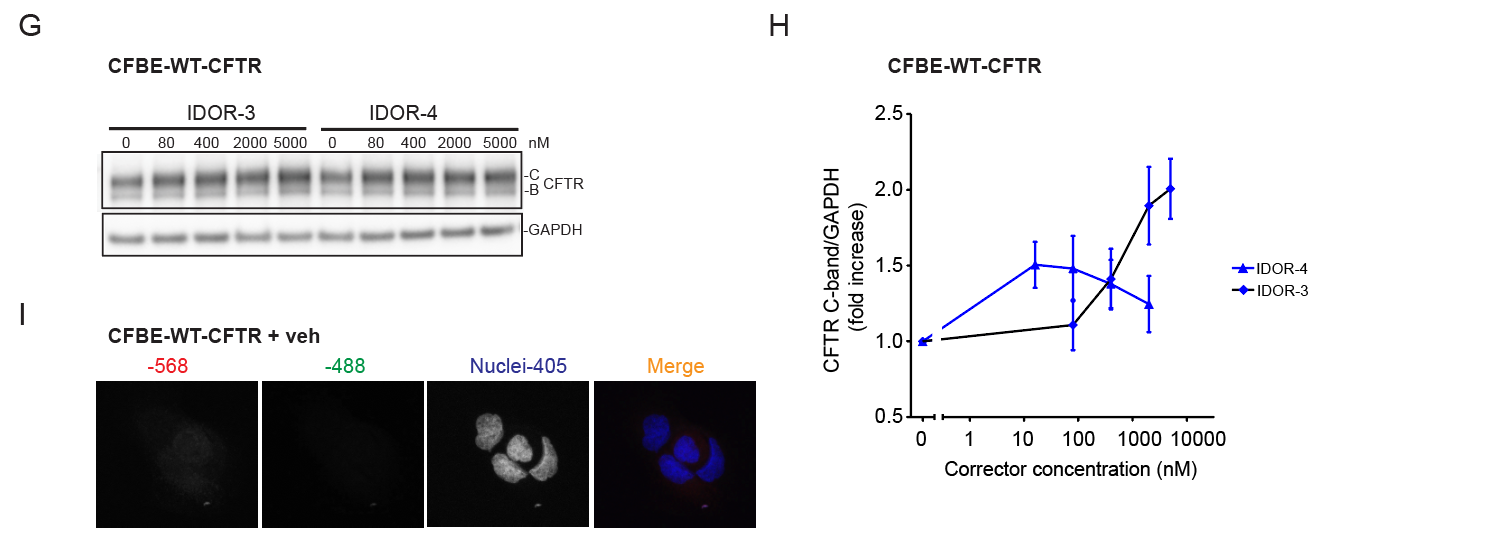

**Fig. S2. Related to Fig. 1: Type-IV correctors restore F508del-CFTR trafficking to the cell surface and behave additively with type-I, type-II, and type-III correctors.** **(A-C)** CFTR surface expression in U2OS-F508del-CFTR cells after over-night treatment with: A) different concentrations of LUM in presence of vehicle or a maximally effective concentration of TEZ (n=3). B) different concentrations of ELX in presence of vehicle or a maximally effective concentration of BAM (n=4). C) different concentrations of IDOR-3 or IDOR-4 in presence of vehicle or a maximally effective concentration of the indicated other correctors (n=3); **(D)** Quantification of CFTR mRNA expression in CFBE41o- cells expressing WT-CFTR (n=2) or F508del-CFTR after 24 h of treatment with DMSO (n=3) or 2000 nM of the indicated correctors (n=2). **(E)** Immunoblot analysis of levels of ubiquitinated proteins, CFTR and GAPDH in CFBE-F508del-CFTR cells after 24 h of treatment with DMSO, MG-132 or the indicated correctors. Representative images (n=3). **(F)** Quantification of ubiquitinated proteins in (E), normalized for GAPDH (n=3). **(G)** Immunoblot analysis of WT-CFTR-expressing CFBE41o- cells treated for 24 h with different concentrations of the indicated correctors and probed for CFTR and GAPDH. Representative images (n=3). **(H)** Quantification of the CFTR C-band intensities in (G) normalized for GAPDH (n=3). **(I)** Representative control immunofluorescence pictures for Figure 1G showing CFBE-WT-CFTR cells treated with DMSO for 24 h and stained for cell nuclei in blue and only secondary antibodies (red and green channels).  Data in (A), (B), (C), (D), (F) and (H) are mean ± SEM of the indicated number of independent experiments.

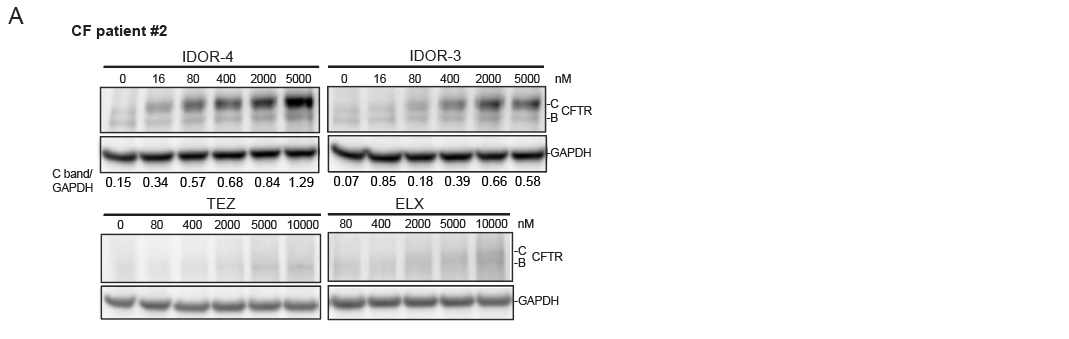

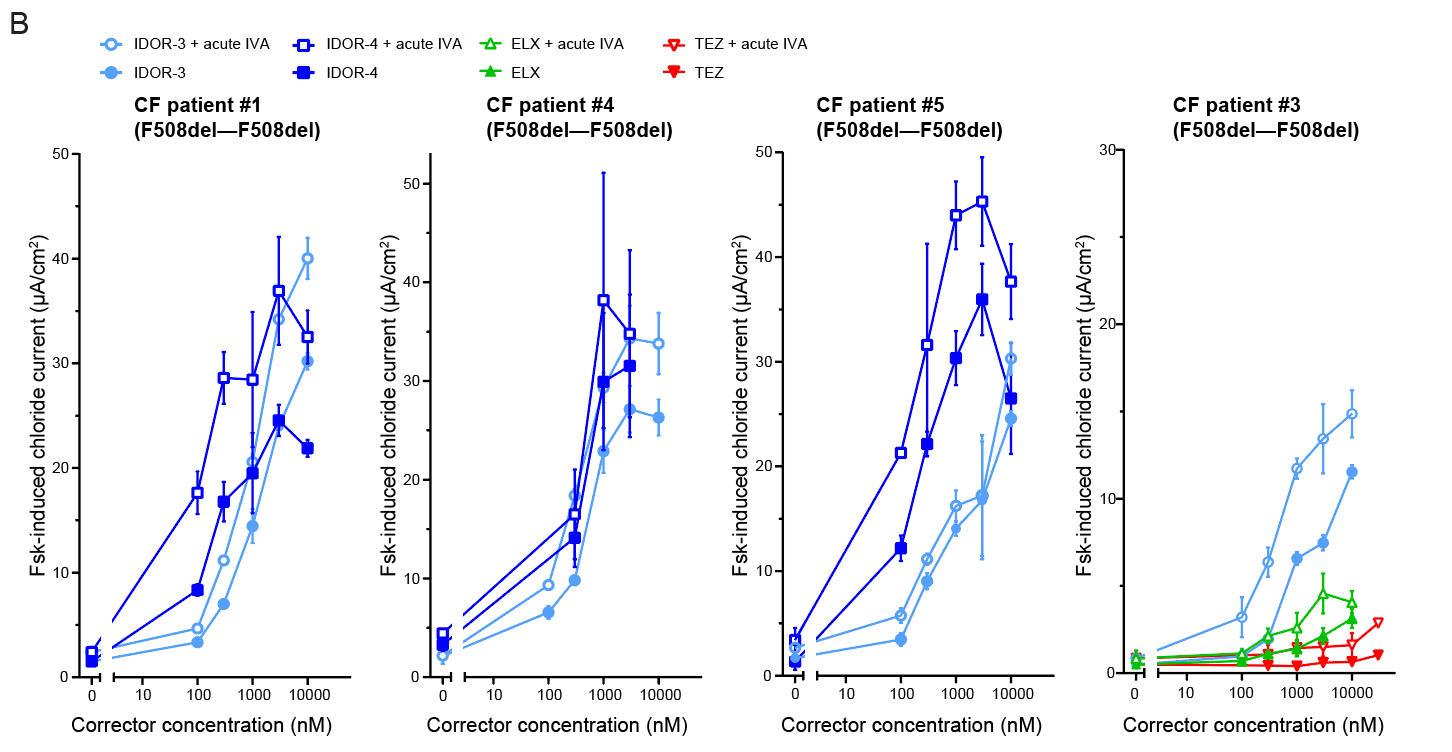

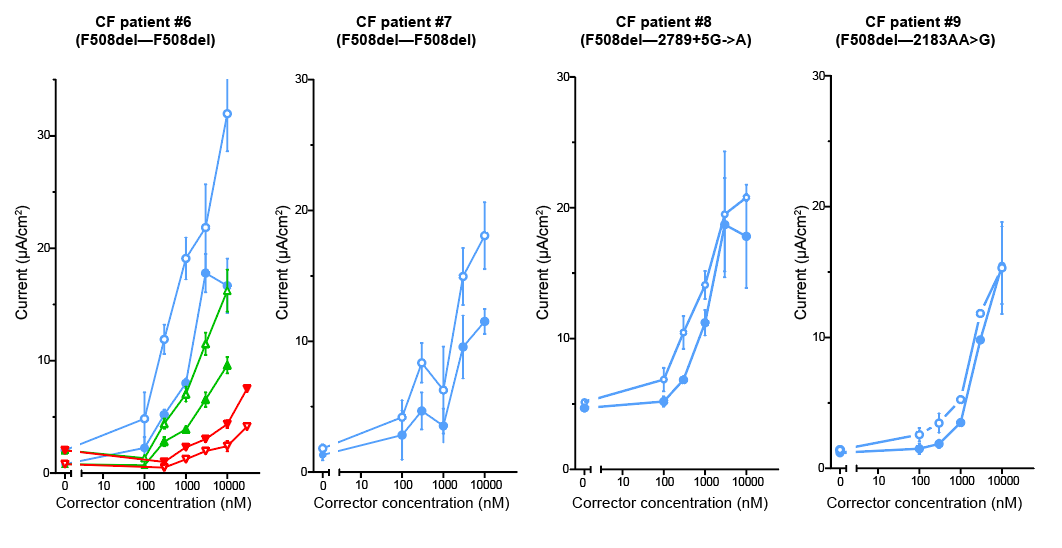

***Fig. S3. Related to Fig. 2: Type-IV correctors rescue F508del-CFTR trafficking and function in reconstituted cystic fibrosis bronchial epithelium.*** ***(A)*** *CFTR expression (immunoblotting) in reconstituted human bronchial epithelium of CF patient #2 (F508del-CFTR homozygous) after a 24-h treatment with the indicated correctors. Representative images (n=2). The CFTR C-band/GAPDH intensity ratio is represented below every lane (for TEZ and ELX correction the ratio could not be calculated because the signal was too weak).* ***(B)*** *Measurement of transepithelial short-circuit currents in Ussing chamber in reconstituted epithelium of 8 CF patients (6 homozygous and two heterozygous for F508del-CFTR), treated over-night with vehicle or IDOR-3, IDOR-4, TEZ or ELX dilution series, in presence or absence of acute 1 µM IVA (at least 2 independent measurements/patient). Data in (B) are mean ± SEM of the indicated number of independent experiments.*

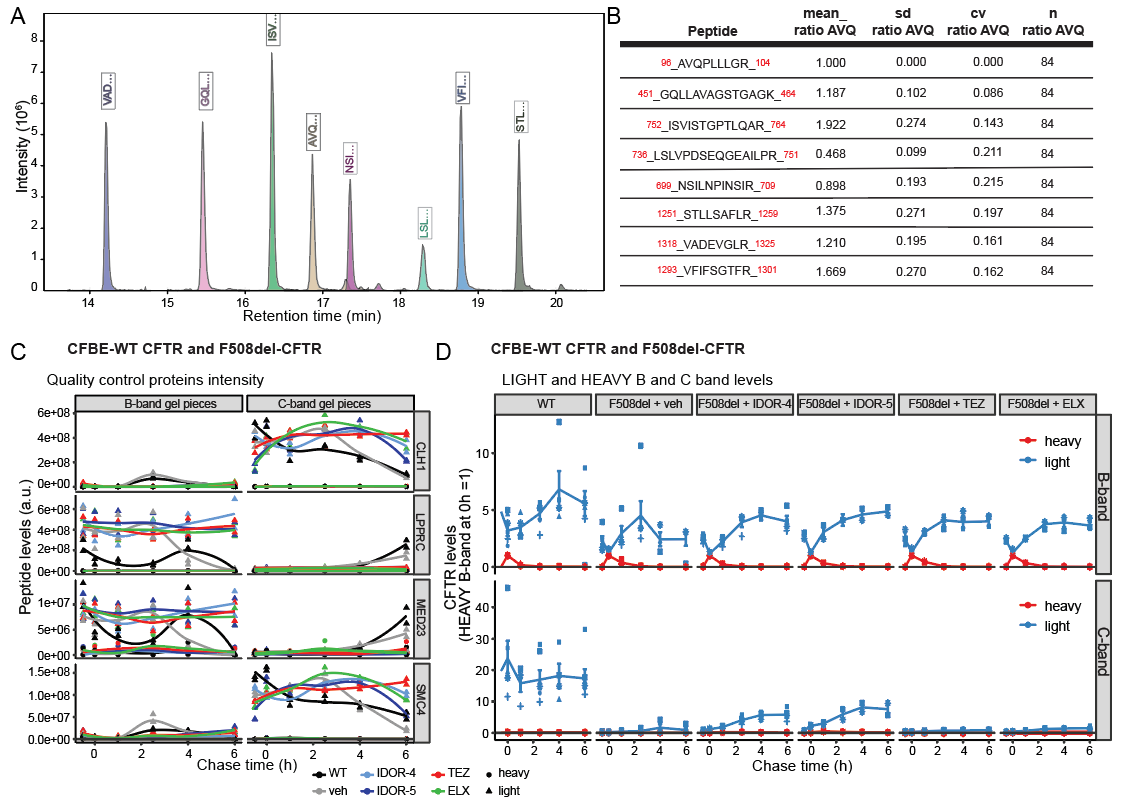

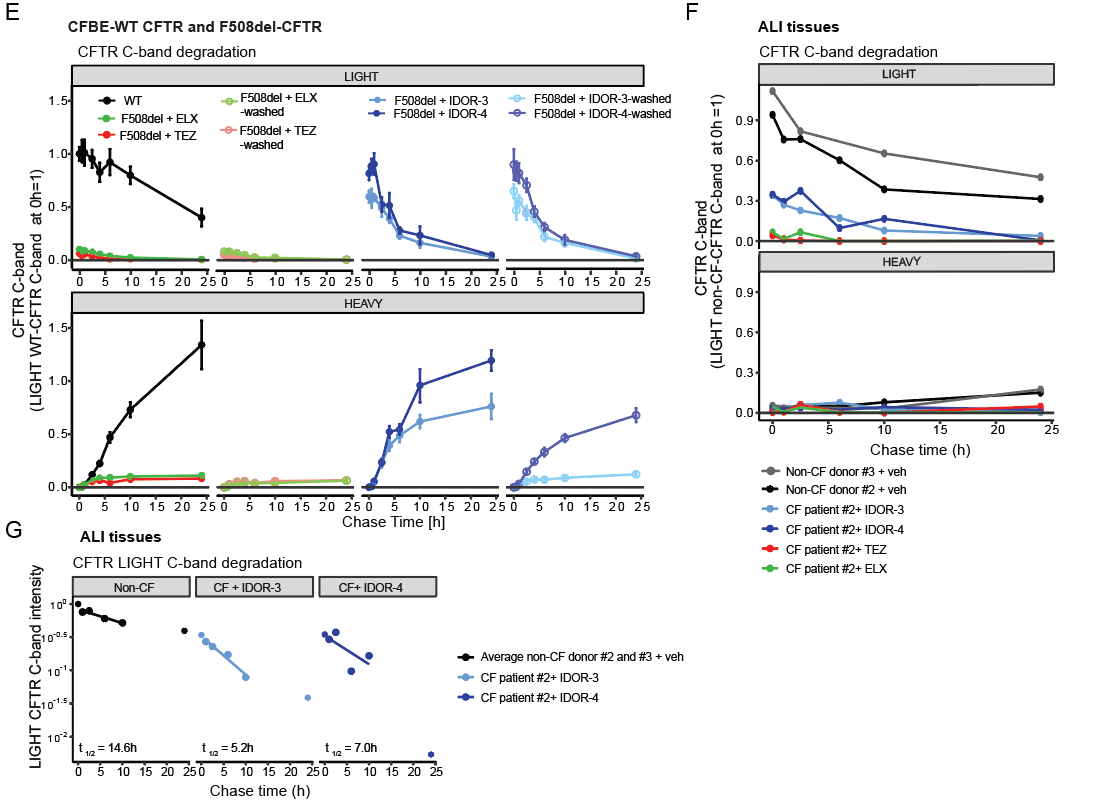

***Fig. S4. Related to Fig. 3: Type-IV correctors restore F508del-CFTR folding efficiency in the endoplasmic reticulum beyond wildtype levels.*** ***(A)*** *Representative chromatogram showing the elution of the 8 peptides quantified for CFTR by mass spectrometry.* ***(B)*** *Representative normalization ratio of each peptide. All peptides are normalized to each other to account for the different response factor using the mean_ratio_AVQ which is the mean ratio of intensity between the respective peptide and the peptide AVQPLLLGR.*  ***(C)*** *Average peptide intensities of heavy-labeled (circles) and light-labelled (triangles) peptides of proteins predicted to co-migrate with CFTR-B band (LPPRC and MED23) or C-band (CLH1 and SMC4). Samples originate from pulse-chase experiments with CFBE cells expressing WT-CFTR or F508del-CFTR, pulse-labeled for 30 min with heavy amino acids and chased with light amino acids for the indicated time periods. During the whole assay F508del-CFTR cells were treated with DMSO (n=5) or 2 µM IDOR-3 (n=4), 2 µM IDOR-4 (n=2), 10 µM TEX or 10 µM ELX (n=3), indicated with different colors.* ***(D)*** *Average peptide intensities of heavy-labeled (red lines) and light-labeled (blue lines) CFTR B- (upper graphs) and C- (lower graphs) bands in CFBE cells treated as described under C). Peptide intensity per time point in each treatment is normalized to maximal intensity of heavy-labeled-B band at 0h chase time.* ***(E)*** *Average peptide intensities of light and heavy CFTR C-band in cells incubated for 24 h with light amino acids in the presence of DMSO (n=4) or 2 µM IDOR-3 or 1 µM IDOR-4 or 10 µM TEZ or 10 µM ELX (n=2 each) and then chased with heavy amino acids. Correctors were either kept during the chase period or thoroughly washed-out before the chase. Per experiment, peptide intensities at the different time points were normalized to the intensity of light C-band of CFTR WT at the beginning of the chase.* ***(F)*** *Peptide intensities of light- and heavy-labeled CFTR C-band in ALI tissues from non-CF donors #2 and #3 and CF patient #2, incubated for 24 h with light amino acids and DMSO or 2 µM IDOR-3, 1 µM IDOR-4, 10 µM TEZ or 10 µM ELX and then chased for the indicated time points with heavy amino acids. Per experiment, peptide intensities were normalized to the maximal intensity of the light-labeled C-band of WT-CFTR at the beginning of the chase (n=1).* ***(G)*** *Calculation of the light-labeled CFTR C-band half-life in ALI tissues from non-CF donors #2 and #3 and CF patient #2 corrected with IDOR-3 and IDOR-4, determined by linear regression of the logarithmic data using time points up to 10 h (n=1). Data in (C), (D) and (E) are mean ± SEM of the indicated number of experiments.*

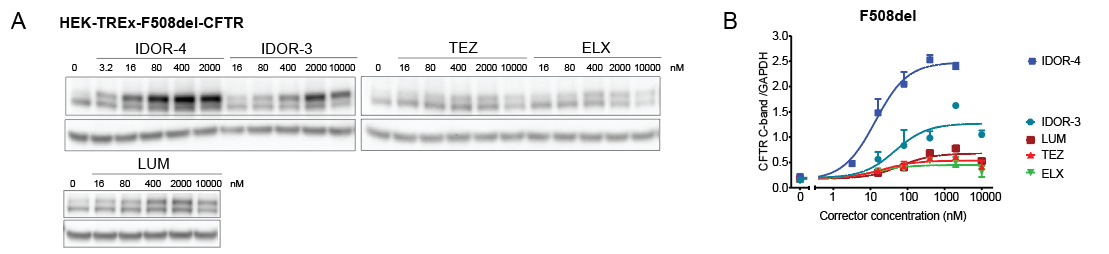

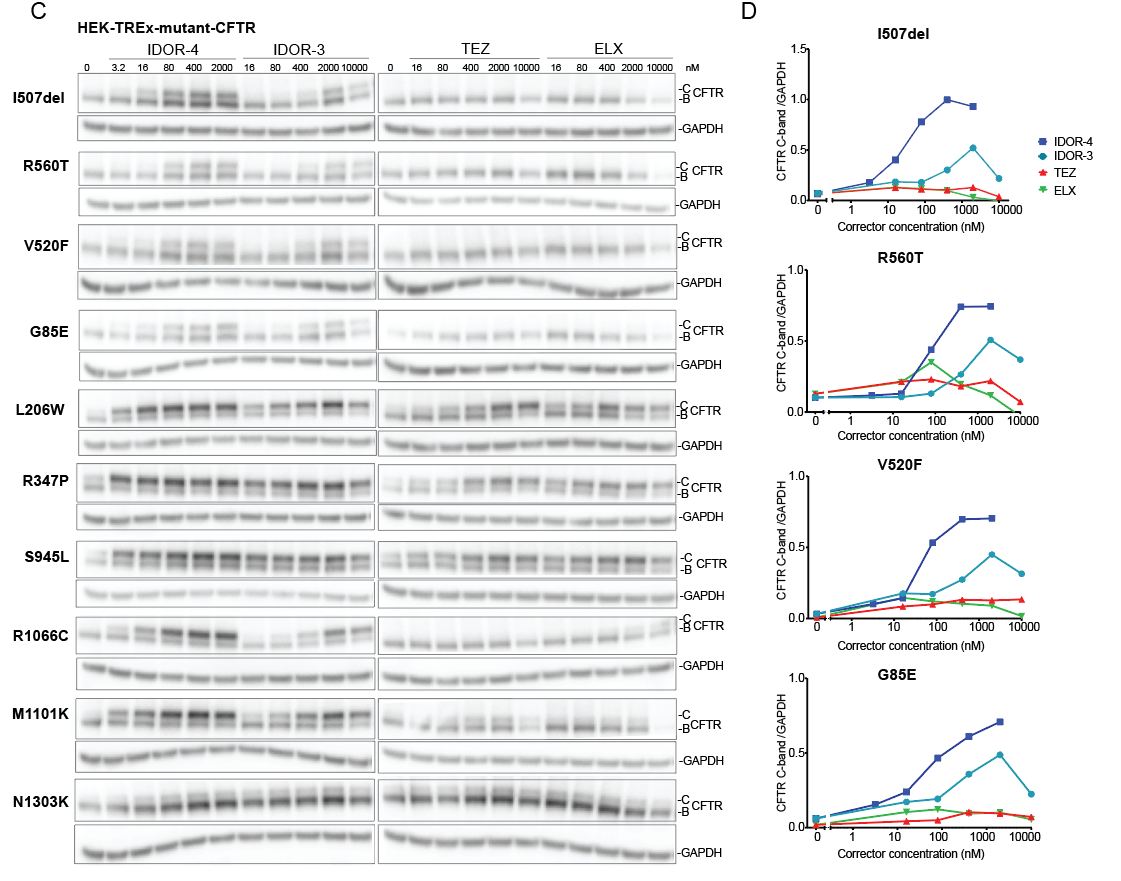

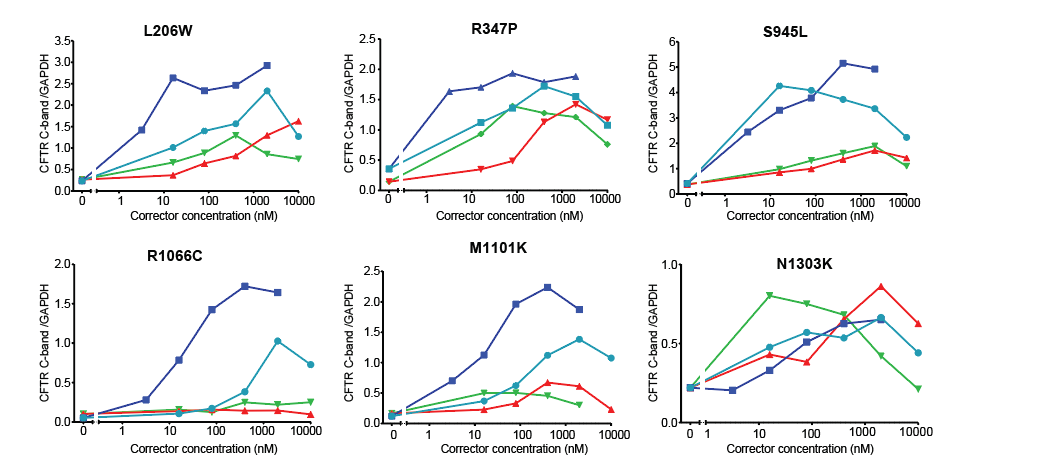

***Fig. S5. Related to Fig. 4 and 6: Type-IV correctors address non-F508del CFTR folding mutations.***

***(A and C)*** *Representative immuno-blot images of CFTR and GAPDH in HEK-TREx cells expressing 11 different CF-causing CFTR folding mutations and treated for 48 h with the indicated corrector concentrations (n=2 for A, n=1 for C). (****B and D)*** *CFTR C-band intensities normalized for GAPDH (n=2 for A, n=1 for C). Data in (B) are means ± SEM of the indicated number of independent experiments. In (D) values are from one experiment.*

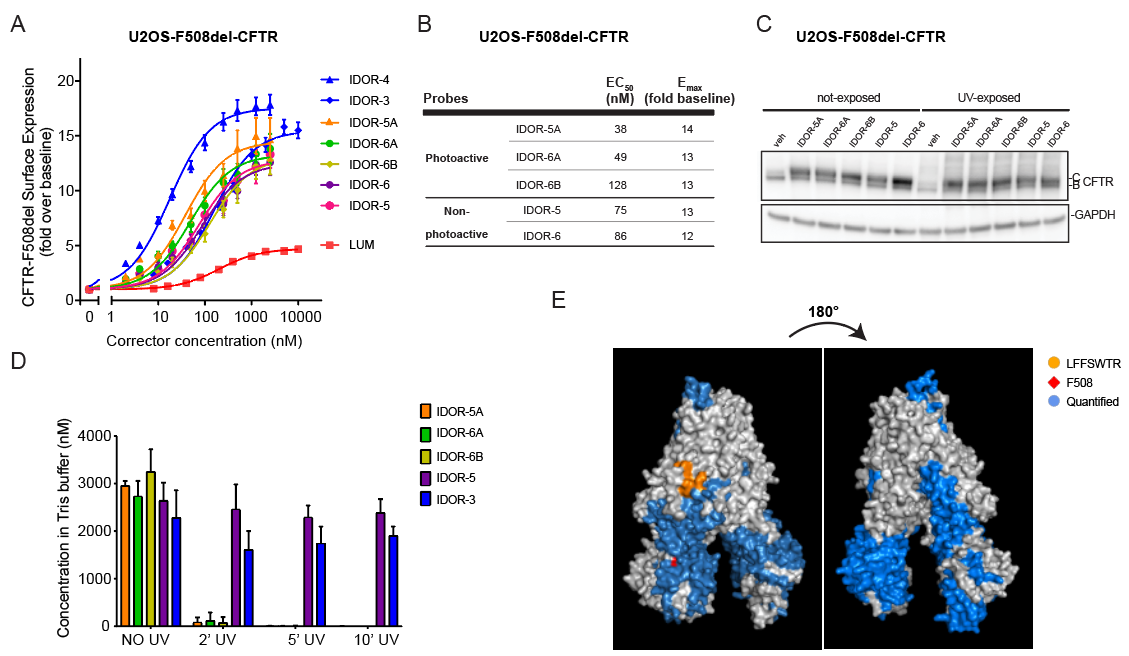

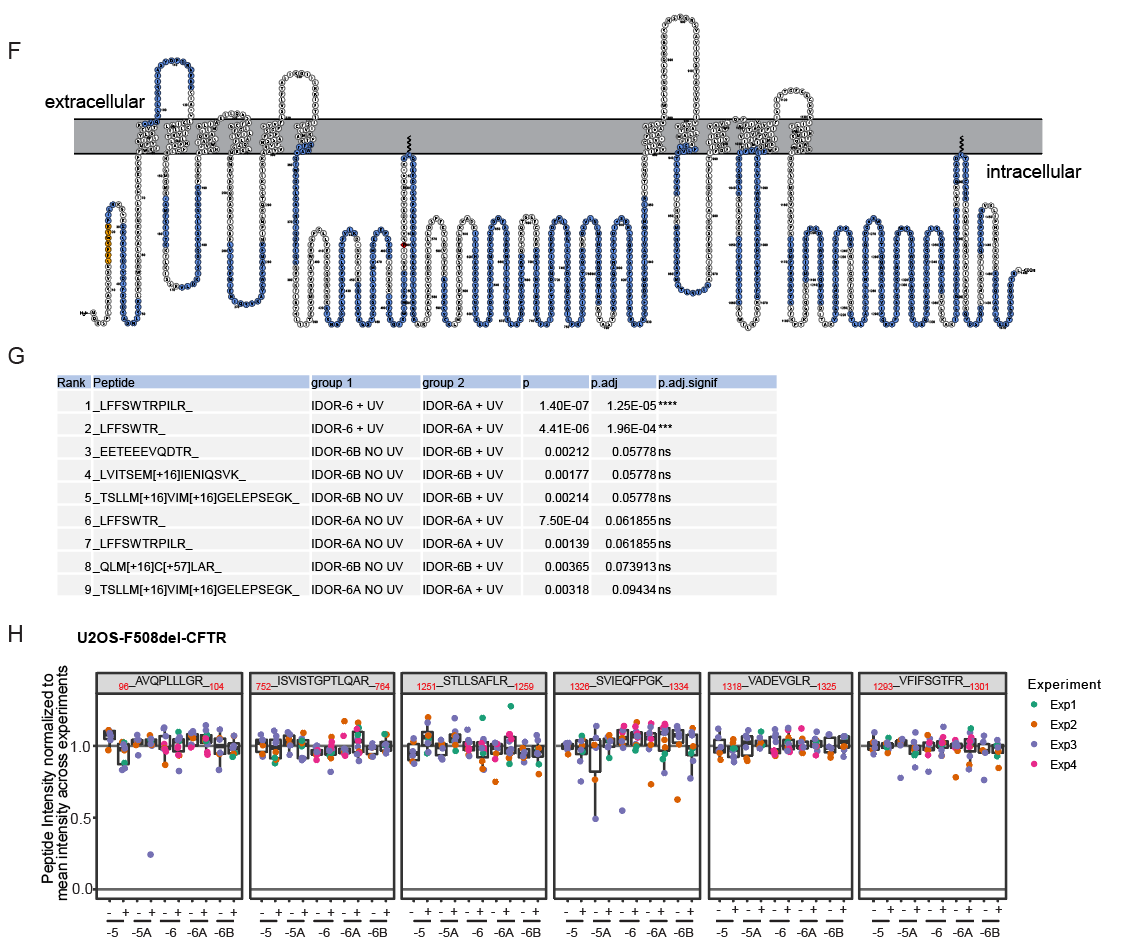
***Fig. S6. Related to Fig. 5: Type-IV correctors bind to the CFTR lasso domain. (A)*** *F508del-CFTR cell surface expression in U2OS cells after overnight treatment with different concentrations of the indicated correctors (n=3 for each probe).* ***(B)*** *EC_50_ and E_max_ values of the three photoactivatable probes and the two parent macrocycles determined in (A).* ***(C)*** *CFTR and GAPDH expression levels (immuno-blotting) in U2OS F508del-CFTR cells treated for 48 h with 2 µM of the indicated probes and exposed or not to UV light for 5 min. Representative images (n=2).* ***(D)*** *Concentration change of the indicated probes diluted in 50 mM Tris buffer with increasing UV exposure times (n=6).* ***(E and F)*** *3D and 2D view of the CFTR protein showing in blue all 73 peptides detected in the crosslinking approach in at least two experiments, corresponding to 53 % of the whole protein. Peptide15-21 is shown in yellow and F508 in red.* ***(G)*** *Table showing the* *first 9 of all identified peptides, ranked from the lowest adjusted p-value after one-sided t-test and multiple-testing correction using Benjamini Hochberg test (see Methods). _15_LFFSWTR_21_ and _15_LFFSWTRPILR_25_ were the only peptides statistically significantly reduced in any comparison and were different for samples treated with IDOR-6A versus the control probe IDOR-6 (after UV exposure). The non-adjusted p-value for these two peptides (without multiple testing correction) showed also a statistically significant difference between IDOR-6A treated samples after and without UV excitation.* ***(H)*** *Abundance of six representative CFTR peptides showing no specific effect after UV exposure or treatment with the six different probes (n=3/4). Data in (A) are mean ± SEM of the indicated number of experiments. Data in (D) are mean ± SD of 6 experimental replica. Scatter plot in (H) results from at least 3independent experiments with colored dots representing replica values. Per experiment, the intensity level of each peptide was normalized to the total CFTR signal in that sample (summed signal of all CFTR peptides).*

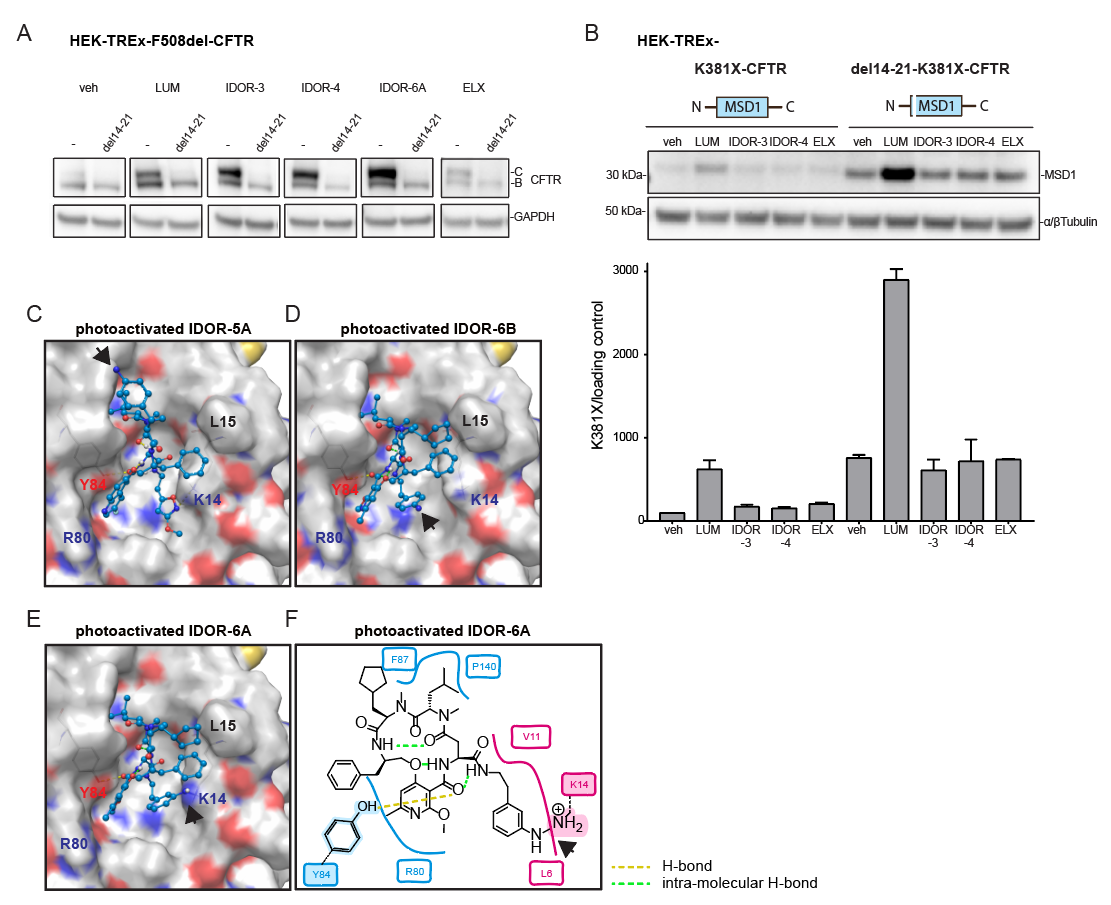

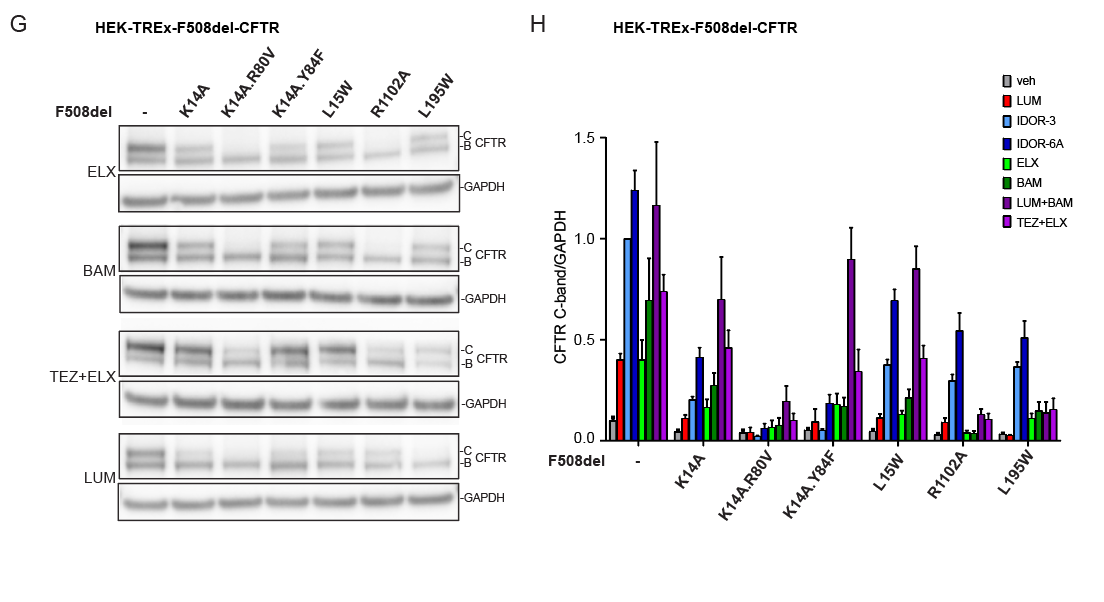
**Fig. S7. Related to Fig. 5: Type-IV correctors bind to the CFTR lasso domain.** **(A)** CFTR and GAPDH expression by immuno-blotting in HEK-TREx cells expressing F508del-CFTR with or without additional deletion of the amino acids 14-21 and treated for 40 h with 2 µM LUM or 2 µM IDOR-3 or 0.4 µM IDOR-4 or 2 µM IDOR-6A or 0.4 µM ELX (n=2). **(B)** Levels of truncated CFTR construct K381X with or without amino acid 14-21 deletion expressed in HEK-TREx cells after 40 h of treatment with vehicle or 2 µM LUM or 2 µM IDOR-3 or 0.4 µM IDOR-4 or 0.4 µM ELX. Detection antibody targets MSD1. Intensities of CFTR fragment bands are shown in the graph below, normalized for the loading controls (n=2). **(C-E)** 3-dimensional views of photo-activated IDOR-5A (C), IDOR-6B (D) and IDOR-6A (E) bound to the cavity as determined by docking studies. Polar amino acid side chains are indicated in blue, red and yellow. H-bonds between the probes and CFTR are indicated as yellow dashed lines. Intramolecular H-bonds are depicted as green dashed lines. Black arrowheads point to the aryl-nitrene (activated intermediate) in each probe. **(F)** 2-dimensional view of IDOR-6A with likely cross-linking of the aryl-meta-nitrene to the K14 amine. **(G)** Site-directed mutagenesis of key amino acids in the proposed type-IV corrector binding site and effect on corrector activities: F508del-CFTR constructs containing one or two additional mutations in the site (K14A, K14A.R80V, K14A.Y84F or L15W) or the ELX binding site (R1102A) or the LUM binding site (L195W) were expressed in HEK-TREx cells then treated for 40 h with 2 µM of each indicated corrector. Representative images (at least 4 independent experiment per mutant and per corrector treatment). **(H)** Quantification of F508del-CFTR C-band intensities in (F) normalized for GAPDH (n=4-10). Data in (B) and (H) are mean ± SEM of the indicated number of experiments.
